# *CASCADES*, a novel *SOX2* super-enhancer associated long noncoding RNA, regulates cancer stem cell specification and differentiation in glioblastoma multiforme

**DOI:** 10.1101/2020.09.05.284349

**Authors:** Uswa Shahzad, Christopher Li, Michael Johnston, Jenny J. Wang, Nesrin Sabha, Frederick S. Varn, Alexandra Riemenschneider, Stacey Krumholtz, Pranathi Meda, Christian A. Smith, Jason Karamchandani, Jonathan K. Watts, Roel G.W. Verhaak, Marco Gallo, James T. Rutka, Sunit Das

## Abstract

Glioblastoma multiforme (GBM) is the most common primary malignant brain tumor in adults, with a median survival of just over one year. The failure of available treatments to achieve remission in patients with GBM has been attributed to the presence of cancer stem cells (CSCs), which are thought to play a central role in tumor development and progression and serve as a treatment-resistant cell repository capable of driving tumor recurrence; in fact, the property of “stemness” itself may be responsible for treatment resistance. In this study, we identify a novel lncRNA, Cancer stem cell associated distal enhancer of *SOX2* (*CASCADES*) that functions as an epigenetic regulator in glioma CSCs (GSCs). *CASCADES* is expressed in IDH-wild type GBM and significantly enriched in GSCs. Knockdown of *CASCADES* in GSCs results in differentiation towards a neuronal lineage in a cell- and cancer-specific manner. Bioinformatics analysis reveals that *CASCADES* functions as a super-enhancer associated lncRNA epigenetic regulator of *SOX2*. Our findings identify *CASCADES* as a critical regulator of stemness in GSCs and represent a novel epigenetic and therapeutic target for disrupting the cancer stem cell compartment in GBM.

## INTRODUCTION

Glioblastoma multiforme (GBM) is the most common primary malignant brain tumor in adults (Aldape et al. 2015; Ostrom et al. 2019). While standard treatment with aggressive multimodal therapy often results in a brief remission, patients uniformly succumb to disease recurrence and progression, with a two-year survival rate of less than 25% (Agnihotri et al. 2013; Nørøxe et al. 2016; Stupp et al. 2005).

This pattern of relapse and progression suggests the presence of a subpopulation of cancer cells within the initial tumor that is able to survive cytotoxic therapies and drive tumor recurrence. Data from multiple studies have supported a hypothesis that the cells responsible for GBM recurrence are cancer stem cells (CSCs) (Singh et al. 2004; Sanai et al. 2005; Gimple et al. 2019; Ma et al. 2018; Lathia and Mack 2015). The CSC hypothesis states that tumor cells are heterogeneous and hierarchically arranged in a manner that recapitulates normal development (Nassar and Blanpain 2016; Kuşoğlu and Biray Avcı 2019; Plaks et al. 2015). Further, the CSC hypothesis posits that, in a manner analogous to normal cell systems, tumor growth is dependent on proliferation of more differentiated cells arising from the CSC population (Plaks et al. 2015; Venere et al. 2011; Beck and Blanpain 2013; Lan et al. 2017). In GBM, glioma CSCs (GSCs) have been found to preferentially survive and be enriched by chemoradiation, suggesting that they may serve as a cell repository for tumor recurrence (Das et al. 2008; Sanai et al. 2005; Venere et al. 2011; Sampetrean and Saya 2013; Bao et al. 2006).

Long noncoding RNAs (lncRNAs) are non-protein coding transcripts longer than 200 nucleotides. LncRNAs have been proposed to regulate processes critical to development, neurobiology, and cancer progression, acting both as transcriptional activators and repressors (Ulitsky and Bartel 2013; Guttman et al. 2011; Khalil et al. 2009; Lam et al. 2014; Rinn and Chang 2012; Kopp and Mendell 2018; Flynn and Chang 2014). As such, lncRNAs have been shown to play an important role in mediating pluripotency, self-renewal, and differentiation programs in embryonic stem cells (ESC) (Guttman et al. 2011; Khalil et al. 2009; Wang et al. 2013). In addition, lncRNAs have been shown to be important in CSC biology (Mineo et al. 2016; Gao et al. 2019; Bhan et al. 2017; Schmitt and Chang 2016).

Among the many mechanisms of action ascribed to lncRNAs, enhancer activity is an important function, primarily because enhancers are capable of driving lineage-specific gene expression (Guttman and Rinn 2012; Rinn and Chang 2012; Li et al. 2016; Lam et al. 2014; Flynn and Chang 2014). A specific class of lncRNAs has emerged that is defined by the presence of active enhancer states (for example, H3K4me1 and H3K27Ac) (Li et al. 2016). These enhancer-associated lncRNAs (lnc-eRNAs) can modulate gene expression both in *cis* and *trans* through chromatin looping, and can dictate temporal and spatial gene regulations during development, differentiation, and cell specification events (Rinn and Chang 2012; Li et al. 2016; Lam et al. 2014).

Super-enhancers (SEs) are groups of putative enhancer clusters in close genomic proximity that are enriched in chromatin features characteristic of enhancers, including transcription factors (TFs), chromatin regulators, co-activators, mediators, RNA polymerase II (RNA Pol II), and enhancer-associated chromatin marks (especially H3K27ac) (Bradner et al. 2017; Huang et al. 2018; Pott and Lieb 2015). SEs are hierarchically organized, containing both hub and non-hub enhancers, based on local chromatin landscape (Huang et al. 2018). The constituent enhancers of SEs can work either independently or as part of a large transcription-regulating machinery to promote incremental expression of their associated genes. SEs tend to regulate genes that are critical in determining cell fate, and thus have cell- and state-specific specialized functions (Bradner et al. 2017; Cheng et al. 2015). It is, therefore, not surprising that SEs have been considered to be essential to maintaining cancer cell identity due to their ability to drive transcription of genes involved in oncogenic processes (Sur and Taipale 2016; Sengupta and George 2017). Super-enhancer RNAs (SE-RNAs) are a class of noncoding RNAs that are transcribed from super-enhancer regions (Chang et al. 2019). Recent studies have suggested that SE-RNAs are functionally important in lineage specification and differentiation processes (Chang et al. 2019; Ounzain et al. 2015).

Given their established role in normal stem cell biology, we hypothesized that lncRNAs could be critical to GSC maintenance in glioblastoma. Here, we report our discovery of *CASCADES*, a novel super-enhancer associated lncRNA that maintains GSC identity through epigenetic regulation of *SOX2*.

## RESULTS

The lncRNAs in close proximity to the transcription factors were selected because a class of *cis*-acting lncRNAs has been shown to directly target their neighboring loci (Kopp and Mendell 2018; Ulitsky and Bartel 2013). Thus, to identify lncRNAs that could modulate GSC identity, we employed an *in silico* “nearest-neighbor” approach for lncRNA candidates in proximity to transcription factors that have been implicated in regulating self-renewal or pluripotency of ESCs, that have been used to successfully reprogram somatic cells into induced pluripotent stem cells (iPSCs), or that have been used to reprogram GSCs, including: POU5F1 (Oct4), Sox2, β-catenin (CTNNB1), Myc, Nanog, Klf4, Zfx, Smad2, Tcf3, Stat3, Fbxo15, GDF3, UTF1, Rex1 (Zfp42), Sall2, and Olig1 (Guttman et al. 2011; Takahashi and Yamanaka 2006; Suvà et al. 2014) **(Figure 1A)**. Using the Ensembl Genome Browser from the European Bioinformatics Institute and the Wellcome Trust Sanger Institute (Flicek et al. 2014), we then evaluated the genomic loci of these transcription factors. We found 112 putative lncRNAs in proximity to these genes of interest (GOI). These lncRNAs were interrogated further using data catalogued in Ensembl, including predicted expression in different cell lines and differential chromatin marks within the genomic loci.

**Figure 1.**
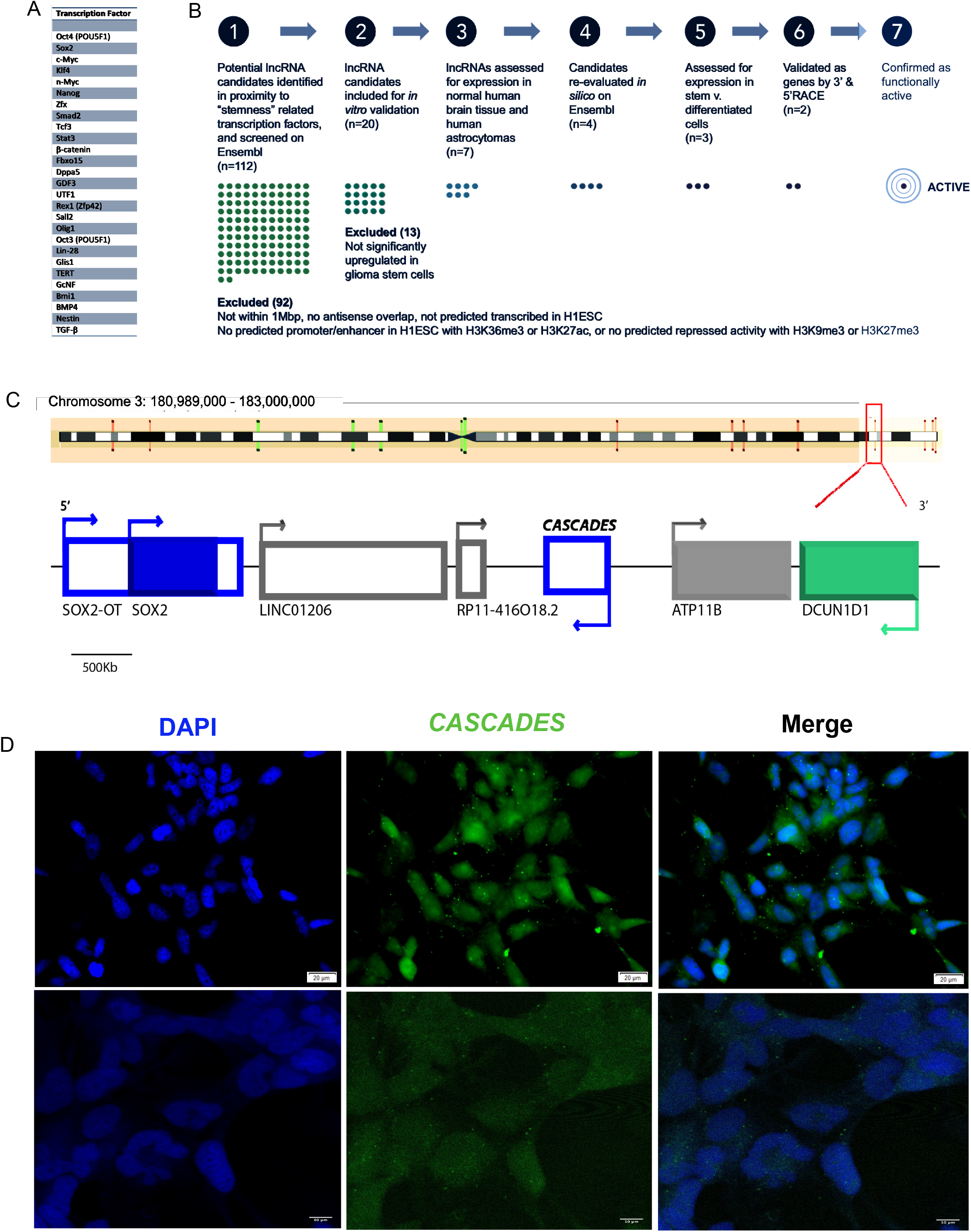
Long noncoding RNA screen to identify novel lncRNA targets relevant to glioma biology. **A**. List of transcription factors used in the *in silico* lncRNA screen. **B**. Pipeline for *in silico* screen to identify novel long noncoding RNAs. **C**. Genomic map depicting the location of *CASCADES* lncRNA in relation to *SOX2*. **D**. RNA *in situ* hybridization of *CASCADES* in glioma stem cell (GliNS1) cell line. Scale bars 20um and 10um.

Following this initial screen, we then interrogated lncRNA candidates using the following criteria: 1) the lncRNA must be within 1 Mbp either upstream or downstream from a GOI; 2) the lncRNA must not have any antisense overlap with the GOI; 3) the lncRNA must be predicted to be transcribed in at least one human embryonic stem cell line (H1ESC); 4) the gene locus containing the lncRNA must have a predicted promoter or enhancer in H1ESC with either H3K36me3 or H3K27ac histone features, or the gene locus must show repressed activity with H3K9me3 or H3K27me3 histone marks. Using this approach, we identified a list of 20 lncRNAs for further study **(Supp Table 1)**. We then validated the expression of the 20 selected lncRNAs in human embryonic stem cells (hESCs), induced pluripotent stem cells (iPSCs), human umbilical vascular endothelial cells (HUVECs), human fetal neural stem cells (HFNS), and GSCs. Based on these findings, we chose to study an intergenic lncRNA found on chromosome 3 at 3q26.33 (gene ID: LINC01995), downstream from *SOX2* and on the opposite strand, which we named <u>C</u>ancer <u>S</u>tem <u>C</u>ell <u>A</u>ssociated Distal Enhancer of *<u>S</u>OX2* (*CASCADES*; **Figure 1B-C**).

Results of RNA *in situ* hybridization and 3’ and 5’ RACE validated *CASCADES* as a transcribed gene **(Figure 1D; Supp Figure 1).**Study of *CASCADES* using the UCSC Genome Browser revealed a gene with two common splice variants that was highly conserved across different species **(Figure 2A)**.

**Figure 2.**
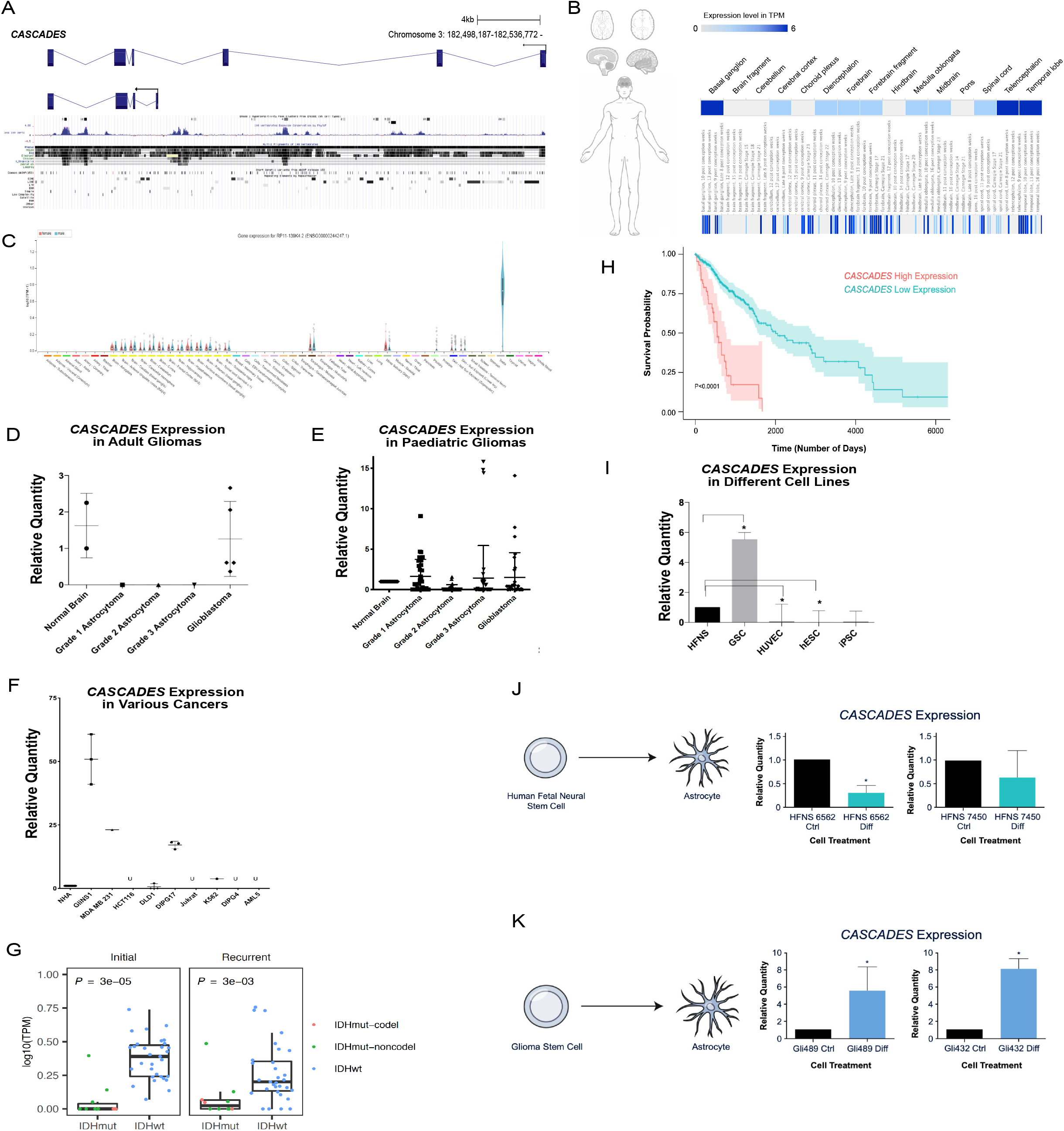
Expression profile of <u>C</u>ancer <u>S</u>tem <u>C</u>ell <u>A</u>ssociated <u>D</u>istal <u>E</u>nhancer of *<u>S</u>OX2 (CASCADES)*. **A**. UCSC Genome Browser tracks show that both of the *CASCADES* transcripts have peaks conserved across many different species. **B & C**. EMBL-EBI Expression Atlas profile of *CASCADES* in the different body tissues and the developing brain. **D**. In adults, *CASCADES* is expressed in the normal brain and grave IV gliomas (glioblastoma) only. **E**. In paediatric population, *CASCADES* is expressed in all grades of gliomas. **F**. *CASCADES* is expressed in gliomas, breast cancer (MDA MB 231), colorectal cancer (DLD1), diffuse intrinsic pontine glioma (DIPG17), and chronic myelogenous leukemia (K592). **G**. RNA-seq of both primary and recurrent gliomas from the Glioma Longitudinal Analysis (GLASS) shows that *CASCADES* is significantly enriched in IDH-wildtype compared to IDH-mutant tumors. **H**. Patient survival data from The Cancer Genome Atlas (TCGA) shows a survival disadvantage for glioma patients with high expression of *CASCADES* (P<0.0001). **I**. *CASCADES* is enriched in glioma stem cells (GSCs) vs. normal fetal neural stem cells (HFNS), human umbilical vein endothelial cells (HUVECs), human embryonic stem cells (hESCs), and induced pluripotent stem cells (iPSCs) (P<0.05). **J**. Upon differentiation of human fetal neural stem cells into astrocytes, *CASCADES* decreased in expression (P<0.05). **K**. When glioma stem cells are forced to differentiate into astrocyte-like cells, *CASCADES* expression is enriched (P<0.05).

Analysis using the EMBL-EBI Expression Atlas suggested that *CASCADES* is expressed mostly in the testes and the developing brain **(Figure 2B-C)**. In adults, *CASCADES* is expressed in normal brain and GBM, with no detectable expression (by qPCR) in low-grade astrocytomas **(Figure 2D)**. In contrast, we found evidence of *CASCADES* expression in all pediatric gliomas, regardless of grade **(Figure 2E)**. In addition to gliomas, we found *CASCADES* expression in breast cancer (MDA MB 231), colorectal cancer (DLD1), diffuse intrinsic pontine glioma (DIPG17), and chronic myelogenous leukemia (K592) **(Figure 2F).**

To clarify the nature of its differential expression in glioma, we interrogated *CASCADES* expression using RNA-sequencing data of both primary and recurrent gliomas from the Glioma Longitudinal Analysis (GLASS) consortium database. These studies showed that in glioma, *CASCADES* is highly expressed in both primary and recurrent gliomas with wild-type isocitrate dehydrogenase (IDH); conversely, we found no significant evidence of expression of *CASCADES* in IDH-mutant gliomas, regardless of 1p and 19q deletion status **(Figure 2G)**. Indeed, the expression of *CASCADES* in pediatric but not adult low-grade gliomas speaks to the frequency of IDH mutation in the latter, and near absence in the former. These data suggest that *CASCADES* expression occurs in a mutually exclusive fashion with IDH mutation in human glioma. Moreover, bioinformatic analysis of the data from the Cancer Genome Atlas revealed that among all glioma patients, the ones who have high expression of *CASCADES* have poor overall survival **(Figure 2H)**.

As lncRNAs have been shown to demonstrate both tissue- and cell-specific patterns of compartmentalization, we then turned to studies to determine if *CASCADES* expression showed cell specificity. We found *CASCADES* to be enriched in GSCs versus normal human fetal neural stem cells **(Figure 2I)**. Interestingly, forced differentiation of normal fetal neural stem cells to astrocytes **(Supp Figure 2A & B)**resulted in a significant decrease in *CASCADES* expression **(Figure 2J)**, while forced differentiation of GSCs to astrocyte-like cells **(Supp Figure 2C & D)**resulted in an unexpected increase in *CASCADES* expression **(Figure 2K)**.

In order to understand the functional role of *CASCADES* in GSCs, we performed siRNA-mediated knockdown *in vitro* using two previously validated primary human GSC lines, GliNS1 and Gli489 (N=3/group; **Supp Figure 2E & F**). *CASCADES* knockdown in GSCs resulted in a significant decrease in *CASCADES* expression, compared to universal (scrambled) control. *CASCADES* knockdown had no adverse effects on cell viability, as assessed using LIVE/DEAD staining and the trypan blue exclusion assay **(Supp Fig 2G - J)**. *CASCADES* knockdown in GSCs did result in a significant decrease in the stemness markers, Nestin **(Figure 3A; Supp Figures 3 & 4)**and Sox2 **(Figure 3B; Supp Figures 3 & 4)**, in comparison to their respective scrambled controls. Interestingly, the expression of the astrocyte marker, GFAP, also significantly decreased after knockdown of *CASCADES* **(Figure 3C; Supp Figures 3 & 4)**. Conversely, CASCADES knockdown resulted in a significant increase in the expression of the neuronal marker, Tuj1, and decrease in the anti-neurogenic transcription factor, Olig2, compared to scrambled control **(Figure 3D-E; Supp Figures 3 & 4)**. Similarly, *CASCADES* knockdown in the human fetal NSC line, HFNS 7450 (N=3/group), resulted in a decrease in Nestin **(Figure 3F; Supp Figures 5 & 6)**and Sox2 **(Figure 3G; Supp Figures 5 & 6)**, compared to scrambled control. However, unlike in GSCs, the expression of GFAP **(Figure 3H; Supp Figures 5 & 6)**, Tuj1 **(Figure 3I; Supp Figures 5 & 6)**, and Olig2 **(Figure 3J; Supp Figure 5)**in human fetal NFCs was unchanged after *CASCADES* knockdown, compared to scrambled controls.

**Figure 3.**
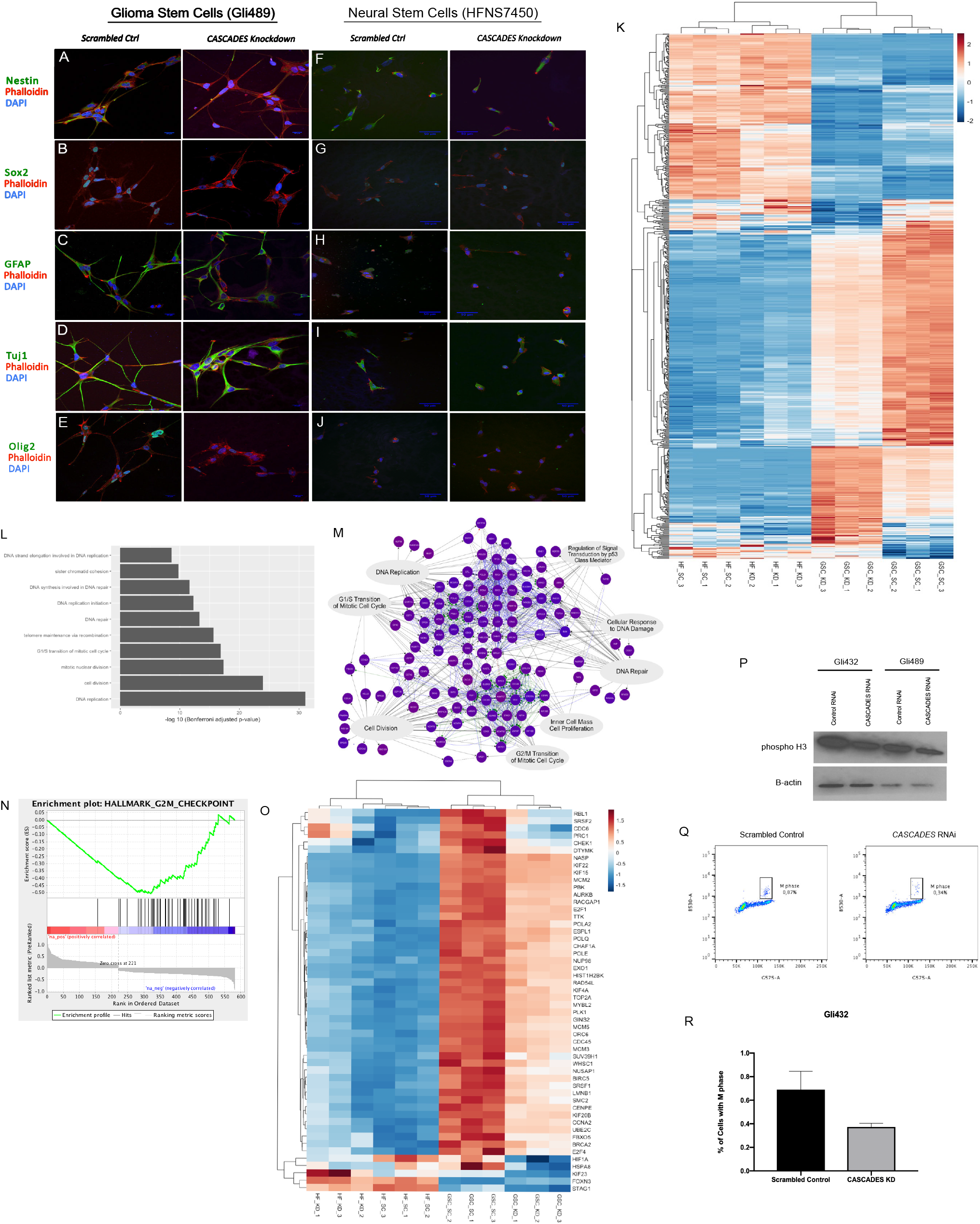
The knockdown of *CASCADES* was performed in glioma stem cells and human fetal neural stem cells (N=3/group) using siRNA. **A, B, & C.**The stem markers, Nestin and Sox2, as well as glial marker GFAP decreased in expression upon the knockdown of *CASCADES* in GSCs. **D & E.**The expression of neural marker, Tuj1, increased upon the knockdown of *CASCADES*, whereas Olig2 decreased. **F & G.**The knockdown of *CASCADES* in HFNS decreased the stem markers, Nestin and Sox2. **H, I, & J.**The expression of glial and neural markers, GFAP, Tuj1, and Olig2, respectively, was unchanged upon knockdown of *CASCADES.* **K**. RNA Sequencing analysis showed differential gene expression upon knockdown of *CASCADES* only in GSCs, whereas this effect was not as pronounced in HFNS. **L**. Gene Ontology of the top 10 pathways perturbed upon the knockdown of *CASCADES*. **M**. Gene Set Enrichment Analysis of the differentially expressed genes upon the knockdown of *CASCADES* in GSCs revealed pathways involved in regulation of cell cycle, mitosis, as well as DNA repair pathways. **N**. Gene Set Enrichment Analysis of the differentially expressed genes upon the knockdown of *CASCADES* in GSCs revealed genes involved in the G2/M check point of the cell cycle. **O**. The knockdown of *CASCADES* results in downregulation of genes involved in the G2/M checkpoint only in GSCs, whereas this effect is unobserved in HFNS. **P**. The expression of cell mitosis marker, phospho H3, decreases upon the knockdown of *CASCADES* in GSCs. **Q & R**. Cell cycle analysis using histone H3 as a marker showed a significant reduction of the cells in M phase of the cell cycle upon knockdown of
*CASCADES* in GSCs, compared to scrambled control (P<0.05).

To determine if *CASCADES* knockdown has a broader effect on gene expression, we then performed RNA-sequencing in GSCs and NSCs before and following siRNA treatment. RNA-seq analysis performed after *CASCADES* knockdown showed differential cellular gene expression in GSCs only, whereas no such effect was observed upon knockdown of *CASCADES* in NSCs **(Figure 3K)**. Gene Ontology (GO) analysis and Gene Set Enrichment Analysis (GSEA) suggested that the effect of CASCADES knockdown in GSCs was predominantly on transcriptional programs involved in DNA replication and control of the cell cycle **(Figure 3L and 3M)**.

GSEA of differentially expressed genes in GSCs and NSCs following *CASCADES* knockdown also showed GSC-specific downregulation of genes involved in the G2/M checkpoint **(Figure 3N-O)**. Indeed, Western blot analysis showed a decrease in phosphorylated H3 expression upon knockdown of *CASCADES* in GSCs **(Figure 3P),**consistent with decreased mitosis. Similarly, cell cycle analysis in GSCs using histone H3 as a marker for mitosis showed a significant reduction in the proportion of cells in the M phase with *CASCADES* knockdown, compared to scrambled control **(Figure 3Q & R)**.

We then sought to determine the mechanism of action of *CASCADES*. First, we performed cellular subfractionation followed by qRT-PCR to identify the subcellular location of the *CASCADES* transcript. While we found evidence of *CASCADES* transcript in the cytoplasm and nucleus, it was found to be mostly chromatin-bound **(Figure 4A; Supp Figure 7A-B)**. Further analysis of *CASCADES* on the Washington University Epigenome Browser showed DNase I hypersensitive sites within the gene body **(Figure 4B)**. Bioinformatic analysis of *CASCADES* revealed a stem cell-specific distal enhancer element for *SOX2* in GSCs. This enhancer element was found within the last intron of *CASCADES*, and was conserved across both splice variants. The presence of H3K4me3 and H3K27ac peaks, flanked by H3K4me1, suggested open chromatin and an active hub enhancer (Ensembl ID: ENSR00001076566). In addition, the second, longer transcript of *CASCADES* contained one other active chromatin site (Ensembl ID: ENSR00000710691) and four other non-hub enhancers (Ensembl IDs: ENSR00000710692, ENSR00000710695, ENSR00000710696, and ENSR00000710698). The neighboring genomic region also contained an enhancer (Ensembl ID: ENSR00000710689), revealing the presence of a super-enhancer within this region. Taken together, this suggests that *CASCADES* is a super-enhancer associated lncRNA. RAMPAGE-Seq data showed the presence of transcriptional start site peaks for *CASCADES* in NSCs but not differentiated neurons **(Figure 4B)**, suggesting a mechanism for its stem cell-specific function.

**Figure 4.**
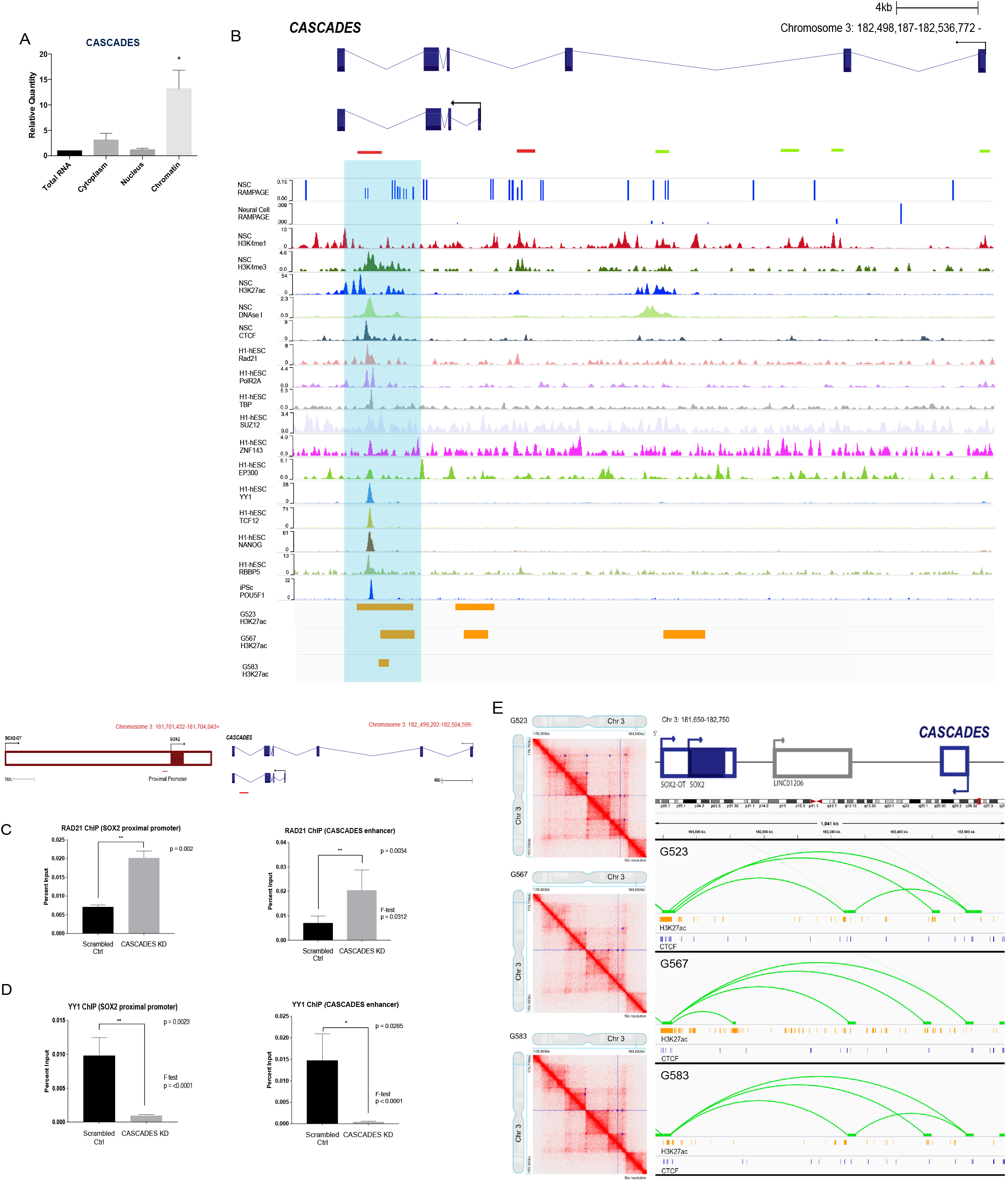
CASCADES is a super-enhancer-associated lncRNA. **A**. Subcellular localization of *CASCADES* showed that it is mostly chromatin-bound. **B**. WashingtonU Epigenome Browser tracks in conjunction with RAMPAGE-Seq data revealed a hub enhancer within the last intron of *CASCADES*, indicated by DNAse I hypersensitive sites, along with open chromatin marks H3K4Me1, H3K4Me3, and H3K27ac. This hub enhancer is conserved across both transcripts of *CASCADES.* The second, longer transcript of *CASCADES* also contains one other open chromatin site and 4 non-hub enhancers. ChIP-Seq of H3K27ac in GSC lines G523, G567, and G583 showed H3K27ac enrichment at the hub enhancer. Red bars indicate open chromatin, whereas green bars indicate non-hub enhancers. Hub enancer is highlighted by blue panel. **C**. ChIP-PCR of Rad21 showed an increased binding of Rad21 at both *SOX2* proximal promoter and the enhancer element found within *CASCADES* **D**. The knockdown of *CASCADES* results in decreased binding of YY1 at the proximal promoter of SOX2 and the *CASCADES* enhancer element. **E**. Hi-C performed in glioma stem cells revealed chromatin loops that occur between the SOX2 proximal promoter and the hub enhancer element found within *CASCADES*. ChIP-Seq of CTCF and H3K27ac in GSC lines also showed enrichment at the loop contact points.

The hub enhancer element also showed binding of transcription factors and proteins that are critical to gene transcription, such as Rad21, YY1, TCF12, as well as core transcription factors like NANOG and POU5F1 (Oct4). Furthermore, ChIP-Seq of H3K27ac in GSC lines (G523, G567, and G583) (Johnston et al. 2019) showed enrichment of H3K27ac at the hub enhancer **(Figure 4B)**. We then sought to test the hypothesis that *CASCADES* could be exerting its effect via chromatin looping events. We first performed chromatin immunoprecipitation (ChIP) of Rad21, a protein mediating promoter-enhancer looping. Rad21-ChIP showed high co-localization of Rad21 at the *CASCADES* enhancer element and *SOX2* proximal promoter. Furthermore, RNAi of *CASCADES* resulted in decreased binding of YY1 and RNA Pol II at the *CASCADES* enhancer element and *SOX2* proximal promoter, and a corresponding increase in Rad21 binding **(Figures 4C & 4D; Supp Figure 8)**. In addition, Chromatin Conformation Capture (Hi-C) of three GSC lines (G523, G567, and G583) (Johnston et al. 2019) confirmed the presence of chromatin loops between the *CASCADES* enhancer element and *SOX2* proximal promoter **(Figure 4E),**further suggesting that chromatin looping is important to its mechanism of action. ChIP-Seq of H3K27ac and CTCF also showed their enrichment at the loop contact points in the GSC lines.

To determine functional differences between the *CASCADES* transcript and the enhancer elements found within *CASCADES*, we designed antisense oligonucleotides (ASOs) targeting various sites across the *CASCADES* gene. We decided to use ASOs to perform knockdown studies, as ASOs are optimal for silencing nuclear, chromatin-associated targets, such as *CASCADES* (Lennox and Behlke 2016). Furthermore, ASOs have been used clinically as central nervous system (CNS) therapeutics (Rinaldi and Wood 2018), making them an attractive option to target a GSC-specific gene like *CASCADES*.

We screened 21 ASOs and identified multiple hits with robust knockdown (at least 70% reduction in mRNA expression) of *CASCADES*. ASOs 8 and 12 were designed proximal to the polyadenylation signal to prevent premature transcriptional termination **(Figure 5A)**(Lee and Mendell 2020; Lai et al. 2020). ASOs 7E and 11E were designed to target the hub enhancer element **(Figure 5B)**. We treated GSCs (Gli432 and Gli489) with each of the four ASOs to silence the expression of *CASCADES* (N=3/group). Consistent with the siRNA data, knockdown of *CASCADES* using ASOs 7E, 8, and 12 resulted in decreased expression of stem cell markers, Nestin and Sox2 **(Figure 5C&D; Supp Fig 9A&B)**. The expression of the neuronal marker, Tuj1, increased upon knockdown using ASOs 11E, 8, and 12 **(Figure 5E; Supp Fig 9C)**, whereas, Olig2 decreased in expression following treatment with all four ASO compounds **(Figure 5F; Supp Fig 9D)**. While the expression of Sox2 reduced upon the knockdown of *CASCADES* using different ASOs, we observed more robust neuronal differentiation using ASOs targeting the 3’ end of the *CASCADES* transcript. These findings show that the *CASCADES* transcript is a crucial component in the mechanism of action for the *SOX2* distal super-enhancer embedded within this genomic region.

**Figure 5.**
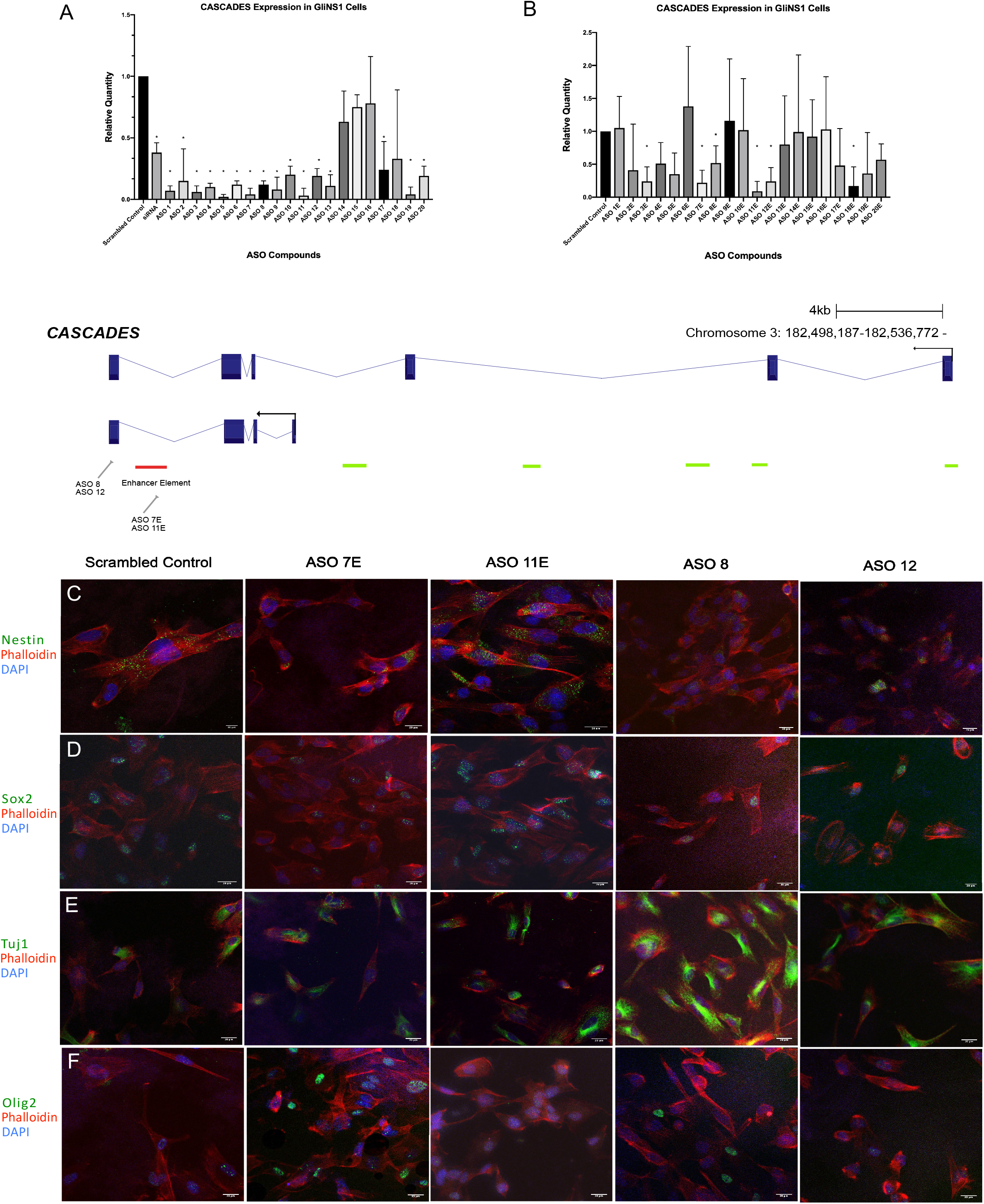
The knockdown of *CASCADES* **was performed in glioma stem cells (N=3/group) using ASOs**. **A**. Antisense oligonucleotides were designed to target various sites across the *CASCADES* transcript. The expression of *CASCADES* was determined using qPCR, and ASO compounds 8 and 12 were selected based upon the extent of knockdown, and the proximity to the poly-A tail. Red bar indicates the hub enhancer, green bars indicate non-hub enhancers. **B**. ASO compounds designed to target the hub enhancer element. Compounds 7E and 11E were selected based on the extent of knockdown. **C & D.**The stem markers, Nestin and Sox2 decreased in expression upon the knockdown of *CASCADES* in GSCs using ASO 7E, ASO 8, and ASO 12. **E**. The expression of neural marker, Tuj1, increased upon the knockdown of CASCADES in ASO 11E, ASO 8, and ASO 12. **F**. The expression of antineurogenic marker Olig2 decreased across all ASO compounds used.

## DISCUSSION

LncRNAs are genetic regulatory elements that, through their role as transcriptional modifiers, can influence global patterns of gene expression in the cells in which they are expressed. As such, lncRNAs have been shown to regulate pluripotency, self-renewal, and differentiation in embryonic stem cells, and have been shown to be critical as mediators of disease progression and metastasis in multiple cancers. Through a combination of *in silico* and *in vitro* studies, we identified a novel lncRNA, *CASCADES*, that regulates GSC identity by acting as a *SOX2* super-enhancer associated lncRNA.

While *CASCADES* is enriched in both neural stem cells and GSCs, our data shows that the knockdown of *CASCADES* induces global effects on cell cycle pathways specifically in GSCs, and promotes GSC differentiation towards a neuronal phenotype. In contrast, no such phenotype is observed upon knockdown of *CASCADES* in neural stem cells. Notably, *CASCADES* expression is limited to IDH-wild type gliomas. Interestingly, GSC differentiation in IDH mutant tumors has been shown to occur toward an astrocytic (AC-like) or oligogodendrocytic (OPC-like) cellular state, but not toward a neuronal (NPC-like) one (Lu et al. 2012; Lathia and Mack 2015; Turcan et al. 2018; Suvà and Tirosh 2020)(Suvà and Tirosh 2020). Conversely, glioma cells that differentiate toward an NPC-like cellular state are found in IDH-wild type glioblastomas. It is intriguing to speculate that *CASCADES*, as a lncRNA specific to IDH-wild type tumors, could account for the specificity of the NPC-like state to IDH-wild type gliomas.

Bioinformatics analysis of *CASCADES* revealed a cell-type specific distal enhancer element for *SOX2* in GSCs, which we determined to be a hub enhancer, and part of a distal super-enhancer of *SOX2*, from which *CASCADES* is transcribed. Super-enhancers are important regulatory elements that generate cell-type specific transcriptional responses. Unsurprisingly, they are central to the maintenance of cancer cell identity because of their role in modulating expression of genes involved in cell growth and lineage specificity (Sengupta and George 2017; Chang et al. 2019). In addition, SEs can promote oncogenic transcription of several growth-related genes, and can be acquired *de novo* during cellular transformation (Sur and Taipale 2016; Sengupta and George 2017). Moreover, SEs show remarkable tumor-type specificity in many different types of cancers, including medulloblastoma (Lin et al. 2016), neuroblastoma (Chipumuro et al. 2014), small-cell lung cancer (Christensen et al. 2014), breast cancer (Wang et al. 2015), esophageal (Jiang et al. 2017) and gastric cancers (Ooi et al. 2016), and melanomas (Zhou et al. 2016; Sengupta and George 2017), thus, making them attractive therapeutic targets in cancer (Sur and Taipale 2016).

It has previously been reported that in mouse embryonic stem cells, super-enhancers associated with *SOX2* contribute to 90% of its expression (Li et al. 2014). Furthermore, in ESCs, co-occupancy of master transcription factors, such as Oct4 (POU5F1), Sox2, and Nanog, along with exceptionally high levels of H3K27ac and DNase I hypersensitivity, is highly representative of super-enhancer activity (Whyte et al. 2013; Hnisz et al. 2013). Our data shows that the *CASCADES* genomic locus, which contains the SE for *SOX2*, has high occupancy of Nanog and POU5F1 at the hub enhancer, along with other important TFs, such as RBBP5, Tcf12, and ZNF143. Additionally, we found that YY1 binds the hub enhancer element as well as the *SOX2* proximal promoter, and that YY1 binding is reduced upon knockdown of *CASCADES*. YY1, also known as Yin Yang 1, has previously been shown to mediate physical enhancer-promoter interactions involving super-enhancers (Weintraub et al. 2017; Thandapani 2019). Our findings support a model in which binding of Rad21 at the *CASCADES* hub enhancer element and *SOX2* proximal promoter suggests chromatin looping, which facilitates RNA Pol II binding at those sites and simultaneous transcription of *CASCADES* and *SOX2*. The *CASCADES* transcript then acts as a "transcription factor-trapper" to entrap YY1 at the proximal promoter of *SOX2* and allow for continued transcription of Sox2 in a positive feedback loop **(Figure 6A)**. Taken together, our data show that *CASCADES* is transcribed from a locus that acts as a super-enhancer for *SOX2*, and that *CASCADES* transcript itself modulates the activity of this SE in a positive feedback loop to promote the expression of *SOX2.*

**Figure 6.**
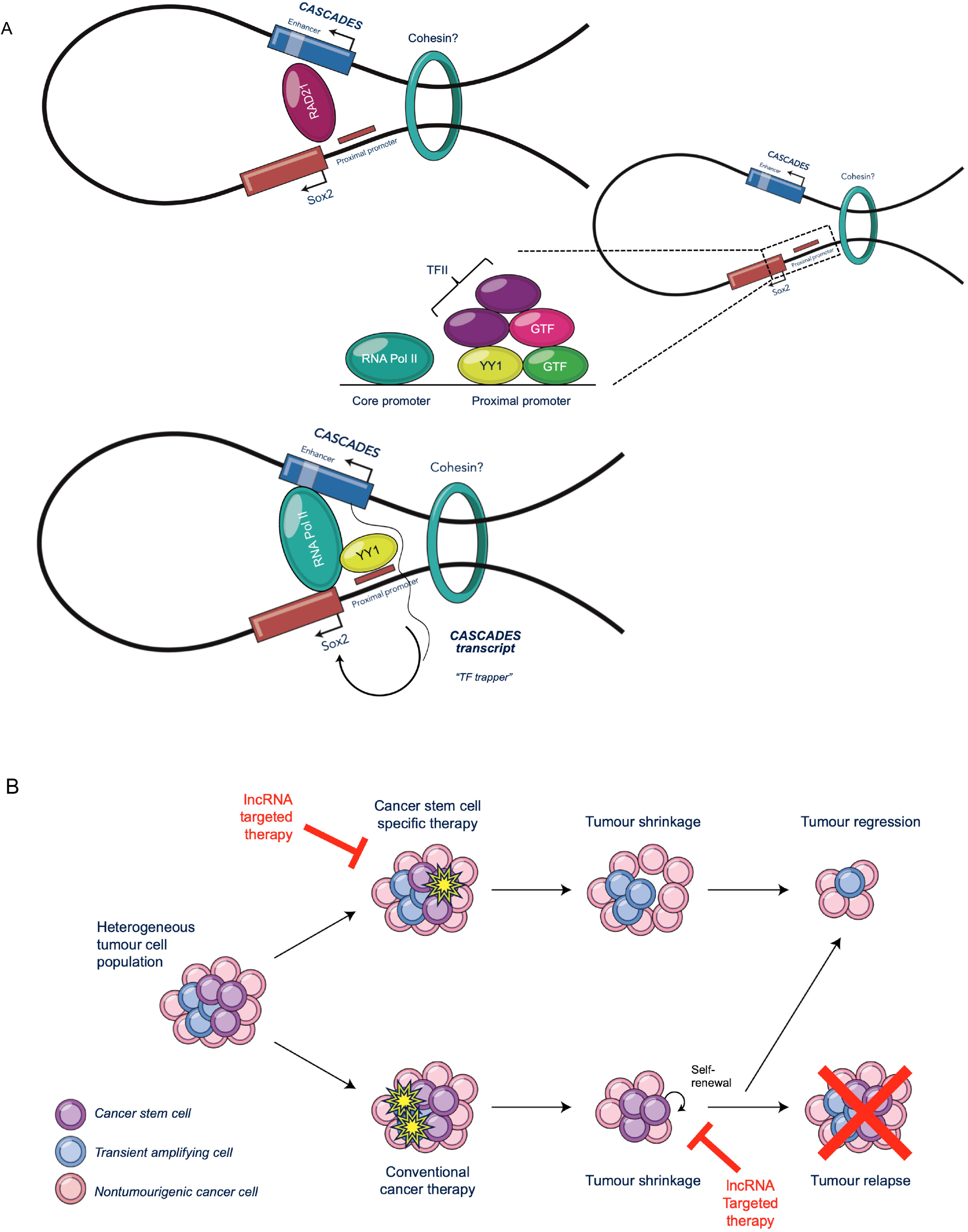
CASCADES is a potential epigenetic target for disrupting the CSC niche in GBM. **A.** The binding of Rad21 at *CASCADES* enhancer and *SOX2* proximal promoter facilitates chromatin looping. This allows for RNA Pol II binding at those sites and for simultaneous transcription of *CASCADES* and *SOX2*. The *CASCADES* transcript acts as a "TF-trapper" and entraps YY1 at the proximal promoter of *SOX2*, and allows for continuous transcription of Sox2 in a positive feedback loop. **B.** The CSC hypothesis posits that within GBM, there is a heterogenous tumor cell population. Conventional cancer therapies target actively dividing cells, whereby CSCs that are capable of becoming quiescent can escape the cytotoxic effects, and can undergo self-renewal and promote tumor relapse. Therapies that specifically target the CSC population can lead to tumor regression. Antisense oligonucleotide therapy targeting a cell- and cancer-specific lncRNA, such as *CASCADES*, can further enhance the effects of anti-CSC thera y, and not only lead to tumor shrinkage, but also prevent tumor relapse.

In summary, *CASCADES* is a novel super-enhancer associated lncRNA that modulates *SOX2* expression and thereby regulates the GSC phenotype, in a manner analogous to other SE associated lncRNAs, such as CCAT1-L(Xiang et al. 2014), *CARMEN* (Ounzain et al. 2015), and LINC01503 (Xie et al. 2018), which have cell type-specific functions in maintaining cell identity and homeostasis. Our data show that the knockdown of *CASCADES* has global effects on cell cycle pathways, specifically in GSCs, and promotes GSC differentiation towards a neuronal phenotype, identifying *CASCADES* as a target to disrupt the GSC niche in glioblastoma and potentially prevent tumor recurrence **(Figure 6B)**.

## CONCLUSION

The importance of the CSC subpopulation in tumor progression and treatment resistance has been demonstrated in multiple hematologic and solid cancers, including glioblastoma. It has been hypothesized that the effective treatment of GBM will require the development of new therapies that are capable of directly targeting GSCs. Our work identifies *CASCADES* as a novel super-enhancer associated lncRNA that modulates *SOX2* expression in GSCs and thereby maintains the cancer stem cell identity in GBM. By targeting a cancer-specific gene, such as *CASCADES*, we can directly target and disrupt the GSC niche in GBM, and potentially prevent tumor recurrence. Further studies involving the use of antisense oligonucleotides targeting *CASCADES* are currently underway to delineate the therapeutic potential of this novel gene in GBM.

## METHODS

All the experiments involving primary human cell lines and human glioma tissue (generously donated by the laboratory of Peter Dirks in the Arthur and Sonia Labatt Brain Tumor Research Centre) were performed in accordance with the research ethics guidelines of the Research Institute at the Hospital for Sick Children, University of Toronto.

### Cell Culture

GliNS1 and Gli489 were cultured and expanded on plates coated with 0.01% poly-L-ornithine (Sigma, Cat # P4957) followed by 1% laminin (Sigma, Cat # L2020) in phosphate buffered saline (PBS) solution. Briefly, the cells were thawed at room temperature, and combined with 10mL of Dulbecco’s Modified Eagle Medium with F12 (DMEM/F12) (Wisent, Cat #219-095-XK) in a 15mL conical tube. By centrifuging at 800rpm for 3mins, the cells were pelleted; the supernatant was aspirated, and the cell pellet was resuspended in 10mL of neural stem cell (NSC) culture media, and plated onto laminin-coated plates. The NSC media was composed of Neurocult NS-A basal medium human (Stem Cell Technologies, Cat #05750) along with L-glutamine (Sigma, Cat #G7513), antibiotic/antimycotic (Sigma, Cat #A5955), bovine serum albumin, epidermal growth factor, basic fibroblast growth factor, B27 supplement, N7 supplement, and heparin. The media was changed every 3-4 days, and the cells were passaged upon reaching ~80% confluence.

### Astrocyte Differentiation

HFNS 6562, HFNS 7450, Gli432, and Gli489 were cultured and expanded on plates coated with 0.01% poly-L-ornithine (Sigma, Cat # P4957) followed by 1% laminin (Sigma, Cat # L2020) in phosphate buffered saline solution, using normal NSC media. The NSC media was composed of Neurocult NS-A basal medium human (Stem Cell Technologies, Cat #05750) along with L-glutamine (Sigma, Cat #G7513), antibiotic/antimycotic (Sigma, Cat #A5955), bovine serum albumin, epidermal growth factor, basic fibroblast growth factor, B27 supplement, N7 supplement, and heparin. The cells were then cultured onto plates coated with Dulbecco’s Modified Eagle Medium with F12 (Wisent, Cat #219-095-XK) containing gelatin (10mg/mL) (Geltrex, Gibco, Cat #A14132-02). Once the cells were 60-70% confluent, they were treated with NSC differentiation media (Neurocult NSA-A basal medium containing L-glutamine, antibiotic/antimycotic, and bovine serum albumin along with 1% fetal bovine serum). Four weeks later, the cells were collected for RNA and protein extraction. The controls were cultured on poly-ornithine and laminin coated plates and were treated with normal neural stem cell media.

### 3’ and 5’ Rapid Amplification of cDNA Ends (RACE)

The 3’RACE (Invitrogen, Cat # 18373-019) and 5’RACE (Invitrogen, Cat #18374-058) were performed using kits and following manufacturer’s instructions. The gene specific primers (GSP) were designed separately using Oligo Primer Analysis software (version 7, Molecular Biology Insights, Colorado Springs, CO). For 3’ RACE, first strand cDNA synthesis was performed in glioma stem cells (GliNS1), human fetal neural stem cells (HFNS 5250), and human umbilical vein endothelial cells (HUVEC), followed by the amplification of target cDNA using first GSP (GSP1). The 1^st^ amplification product was purified using PureLink PCR Purification Kit (Invitrogen, Cat #K3100), following manufacturer’s instructions, and was visualized on a gel. The purified 1^st^ amplification product was then amplified using a nested primer (GSP2), and the 2^nd^ amplification product was purified again, visualized on a DNA gel, and subsequently cloned. On the other hand, for 5’RACE first strand cDNA for 5’RACE was synthesized from GliNS1 using GSP1. The cDNA product was then purified, and a dC-tail was added to this product. Then nested amplification was performed using GSP2. An additional nested amplification was performed, if needed, using GSP3. The DNA products were purified after each amplification, and were visualized on DNA gels. The final nested amplification product was used for cloning.

### Cloning

The 3’ & 5’RACE products were cloned using TOPO TA Cloning Kit (Invitrogen, Cat #K4530-20), following the manufacturer’s instructions. Briefly, the products were ligated using the TOPO vector and incubated at room temperature for 15 minutes. The ligated products were then transformed into *E. coli* through electroporation and plated on kanamycin coated agar plates, followed by incubation at 37°C overnight. Then 3-5 colonies were incubated in lysogeny broth (LB) medium, along with ampicillin, at 37°C overnight. The plasmids were isolated using Plasmid Mini Prep Kit (Frogga Bio, Cat #PD300), and products were incubated with EcoRI for restriction digest and visualized on a DNA gel for validation.

### siRNA-Mediated Knockdown of lncRNAs

The cells were cultured in 6-well plates, using normal NSC culture medium. Twenty-four hours before siRNA knockdown was performed, the cells were incubated with NSC culture medium without antibiotic/antimycotic. The siRNAs were designed using Custom RNAi Design Tool by Integrated DNA Technologies (IDT) (Coralville, Iowa), and were resuspended in Tris-EDTA (TE) buffer to a final concentration of 20μM. For each well, 2μL of siRNA was combined with 138μL of Opti-Mem Reduced Serum Medium with GlutaMax Supplement (Life Technologies, Cat #51985-034) and incubated at room temperature for 5 minutes. Meanwhile, 16.5μL of Oligofectamine Transfection Reagent (Life Technologies, Cat #12252-011) was combined with 43.5μL of Opti-Mem. The diluted siRNA + Opti-Mem was then combined with Oligofectamine + Opti-Mem, and incubated at room temperature for 15 minutes to allow complexes to form. The GliNS1 cells were washed once with Opti-Mem, and 1.3mL of Opti-Mem was added to each well, followed by addition of 200μL of siRNA+Oligofectamine complex. The cells were then incubated at 37°C for 4 hours, and the media was replaced by NSC culture medium without antibiotics/antimycotics. The RNA was extracted 72 hours later for analysis by qRT-PCR.

### ASO design and synthesis

ASOs were designed using the LNCASO webserver (https://iomics.ugent.be/lncaso). ASOs were synthesized as fully PS-modified 5-10-5 MOE gapmers (i.e. 5 nucleotides of 2’-O-methoxyethyl-RNA (MOE), 10 DNA nucleotides, 5 MOE nucleotides) using standard phosphoramidite methods on a Dr.Oligo 48 synthesizer (Biolytic, Freemont, CA). Phosphoramidites and standard reagents were purchased from ChemGenes (Wilmington, MA). Coupling time for MOE nucleotides were extended to 2 min. Oligonucleotides were cleaved and deprotected in concentrated aqueous ammonia at 55°C for 16 h. ASOs were characterized by LC-MS analysis (Table x) using an Agilent Q-TOF instrument and were desalted using Amicon ultrafiltration columns (3-kDa cutoff). The cells were transfected using ASOs following the similar protocol as siRNA-mediated knockdowns.

### Immunocytochemistry

Briefly, the cells were washed with PBS, and fixed with 4% paraformaldehyde (PFA) at room temperature for 10 minutes. They were then permeabilized using 0.3% Triton-X100 in PBS at room temperature for 5 minutes, and were washed twice by PBS. The fixed cells were then blocked with 5% BSA in PBS for 1 hour at room temperature, and were then stained using antibodies against Nestin (Millipore, Cat# MAB5326) and Sox2 (Abcam, Cat# ab97959) (stem markers), glial fibrillary acidic protein (GFAP) (Santa-Cruz, Cat# sc-6171) (astrocyte marker), beta-tubulin III (Tuj1) (Abcam, Cat# ab78078), and Olig2 (R&D, Cat# AF2418) (neuron markers). After incubating with primary antibodies at room temperature for 2 hours, the cells were then washed twice with high salt PBS, and once again with regular PBS. They were then incubated with appropriate Alexa fluor secondary antibodies, along with phalloidin, at room temperature for 1 hour. The cells were then washed as described above, and mounted with DAPI Vectashield media (Vectashield, Cat # H-1500). The slides were then visualized using confocal microscopy (N=3 slides/group, and at least 3-6 different fields of view were imaged). Finally, the number of “positive” cells as well as all cells on a field was counted using ImageJ software and plotted as a percentage of positive cells/total cells.

### Western blot analysis

Total cell lysates were prepared by harvesting cells in RIPA lysis buffer (Sigma-Aldrich, St, Louis, MO) with a protease inhibitor cocktail (Roche Diagnostics, Indianapolis, IN). Protein concentration was determined using the Pierce BCA Protein Assay Kit (Thermo Scientific, Rockford, IL). Protein extracts were mixed with 6X SDS sample buffer (Tris pH 6.8, 1.7% SDS, glycerol and β-mercaptoethanol), and the cell lysates were resolved on 12% SDS-polyacrylamide gels of 1.5 mm thickness. Proteins were then transferred onto polyvinylidene Fluoride Transfer Membranes (Pall Corporation, Pensacola, FL), and subsequently blocked with 5% skim milk in TBST (20mM Tris aminomethane, 150mM NaCl and 0.05% Tween 20; pH 7.4) for 1 hr at room temperature. The membranes were incubated overnight at 4 °C with primary antibodies, then at room temperature for 1 hr with the secondary antibodies, either horseradish peroxidase conjugated goat anti-rabbit or anti-mouse immunoglobulin G antibody (1:5000; Cell Signaling Technology), and bound primary antibodies were visualized using Western Lightning Plus-ECL (PerkinElmer Inc., Waltham, MA). The primary antibodies used in this study were as follows: Nestin (Millipore, Cat# MAB5326), Sox2 (Abcam, Cat# ab97959), glial fibrillary acidic protein (GFAP) (Santa-Cruz, Cat# sc-6171), beta-tubulin III (Tuj1) (Abcam, Cat# ab78078), and Olig2 (R&D, Cat# AF2418), phospho H3 (pSer^10^) (Sigma 9710S).

### LIVE/DEAD Staining

Cell viability was assessed using LIVE/DEAD Cell Imaging Kit (488/570) (Thermo Fisher Scientific, Cat# R37601). Following manufacturer’s protocol, the cells were kept in their original culture vessels. The contents of LIVE green vial were combined with DEAD red vial to create a 2X stock. Then equal volumes of 2X stock were added to each well, and incubated for 15 minutes at room temperature. The cells were then immediately visualized on fluorescence microscope by a blinded observer. The images obtained were then given to another blinded observer, who provided the final count of LIVE (green) and DEAD (red) cells for each sample.

### Cellular Subfractionation

The GliNS1 cells were collected and RNA was extracted, and was partitioned into chromosomal, nuclear, and cystoplasmic fractions, as follows. The cells were washed twice with 10mL PBS−/− and detached using 1mL of accutase, and incubate at 37C for 5 mins. GliNS1 were then collected with DMEM/F12, and added to 10mL conical (Falcon) tube, and pelleted by centrifugation at 14,000rpm for 4 mins. The pellet was then resuspended in cold 1xPBS/1mM EDTA and transferred to a sterile epitube, and centrifuged again to pellet cells at 14,000rpm for 4 mins at 4C. The supernatant was discarded, and the wash was repeated once. The pellet was then resuspended in ice-cold 100uL lysis buffer (10mM Tris-HCl, pH 7.5; 150mM NaCl; 0.15% Nonidet P-40), and lysed for 5 mins on ice. The lysate was layered on top of 2.5 volumes of chilled sucrose cushion (24% sucrose in 10mM Tris-HCl and 150mM NaCl), and centrifuged at 14,000rpm for 10 min at 4C. The supernatant, which contained the cytoplasmic fraction, was treated with proteinase K for 1hr at 37C, followed by RNA extraction and purification using QIAGEN RNeasy Kit. The pellet was retained on ice until ready for further subfractionation.

The pellet was washed once with ice-cold 1xPBX/1mM EDTA, and resuspended in pre-chilled glycerol buffer 50uL (20mM Tris-HCl, pH 7.9, 75mM NaCl, 0.5mM EDTA, 0.85mM DTT, 0.125mM PMSF, 50% glycerol), by gently flicking tube. Then the equal amount (50uL) of cold nuclei lysis buffer (10mM HEPES, pH 7.6, 1mM DTT, 7.5mM MgCl2, 0.2mM EDTA, 0.3M NaCl, 1M Urea, 1% NP-40) was added, and vortexed for 2×2 sec, incubated for 2 min on ice, and then centrifuged for 2 min at 14,000rpm at 4C. The supernatant, which contained soluble nuclear fraction, was treated with proteinase K for 1hr at 37C, and the RNA was extracted using the QIAGEN RNeasy Kit. The pellet was rinsed with cold 1XPBS/1mM EDTA, and dissolve in TRIzol (Invitrogen). The RNA was extracted using TRIzol, and then purified using the QIAGEN RNeasy Kit. To validate appropriate partitioning of nuclear, chromatin-bound, and cytoplasmic RNA, the expression of MALAT1 and Xist was evaluated.

### Chromatin Immunoprecipitation (ChIP) PCR

For chromatin immunoprecipitation, the EZ-Magna ChIP Kit (Millipore, Cat# 17-408) was used, following the manufacturer’s instructions. Briefly, ~20 million cells per ChIP reaction were grown according to the cell culture conditions discussed above. On the day of ChIP, the cells were fixed on culture dish with fresh formaldehyde added directly to the growth media to a final concentration of 1%, and incubated at room temperature for 10 minutes to allow for crosslinking. The media was removed, and 2mL of 10x glycine was added to each dish to quench the unreacted formaldehyde and incubated at room temperature for 5 minutes. The media was aspirated and the cells were washed with cold 1xPBS twice, followed by addition of 2mL of cold 1xPBS/protease inhibitor (PI). The cells were scraped and centrifuged at 800 x g at 4°C for 5 minutes. The supernatant was removed, and the cell pellet was resuspended in 0.5mL of cell lysis buffer/PI and incubated on ice for 15 minutes with brief vortexing. The cell suspension was again centrifuged at 800 x g at 4°C for 5 minutes to remove the cellular fraction. The nuclei were then lysed with nuclear lysis buffer/PI and sonicated using Diagenode Bioruptor with 30 sec on/off cycle for 2 hours at 4°C. The chromatin was sheared to produce fragments of ~200bp in size. The sheared chromatin was then aliquoted, diluted, and 1% volume was collected to serve as an “input” control. To the remaining sheared chromatin, 20uL of fully suspended protein A magnetic beads, along with 5ug of immunoprecipitating antibody for Rad21 (Abcam, Cat# ab992) or YY1 (Santa Cruz, sc-1703X) was added. For the positive control, an anti-acetyl histone H3 antibody, and for negative control, the normal rabbit IgG was added and the reactions were incubated for 4 hours at 4°C with rotation. The protein A bead-antibody/chromatin complex was then washed with the following ice cold wash buffers, respectively: low salt immune complex, high salt immune complex, LiCl immune complex, and TE buffer. For each tube, 100uL of ChIP elution buffer with 1uL of proteinase K was added, and incubated at 62°C for 2 hours with shaking, followed by incubation at 95°C for 10 minutes. The samples were cooled to room temperature, and the DNA was purified using spin columns, and was used subsequently for real-time PCR.

### *RNA fluorescence* in situ hybridization

Cells were grown on glass cover slips in 6-well plates until 70% confluent. Cells were then fixed in 4% cold PFA for 20 minutes, washed, and hydrated. The cells were hybridized with the human *CASCADES* RNA probe in accordance with the manufacturer’s protocols (Advanced Cell Diagnostics, Newark, CA). In brief, cells were pretreated with hydrogen peroxide, permeabilized by incubating with Protease III (1:15) for 10 mins. After hybridization, a fluorescent kit (version 2) was used to amplify the mRNA signal, and TSA Plus Fluorescein, fluorescent signal was detected using a microscope slide scanner (Olympus) and Confocal microscopy. Representative images were prepared in Cell image software (Olympus) and ImageJ.

### Flow Cytometry Cell Cycle Analysis

Cells were collected by centrifugation at 400 xg for 5 minutes at 4°C. The cells were then spun and re-suspended in 50 μl staining medium (HBSS with 2% FBS and azide) and then added to a conical polypropylene tube containing 1 ml of ice-cold 80% ethanol. The cells were then vortexed and fixed overnight at 4°C. The fixed cells were then collected by centrifugation at 400 xg for 5 minutes at 4°C and washed twice with 1X PBS, then once with staining medium (SM). They were then stained with anti-Histone H3 (Phospho-Histone H3 (Ser28) Monoclonal Antibody (HTA28), Alexa Fluor 488, eBioscience) (1:500) for 20 minutes at room temperature, then washed with SM. The pellet was re-suspended in 500 μl 2 mg/ml RNase A solution, and incubated for 5 minutes at room temperature. Then 500 μl 0.1 mg/ml PI solution was added and vortexed to mix. The pellet was incubated for 30 minutes at room temperature, protected from light, and filtered through Nitex into FACS tubes. Finally, the cells were run through a BD analyzer, and the results were analyzed using FlowJo software.

### Bioinformatics Analysis

LncRNA expression data was obtained from the Wellcome Trust Ensembl Genome Browser (Flicek et al. 2014). *CASCADES* (LINC01995) tissue expression data was collected from EMBL-EBI Expression Atlas (https://www.ebi.ac.uk/gxa/home). The splice variants were identified using NCBI (https://www.ncbi.nlm.nih.gov/gene/?term=LINC01995) and UCSC Genome Browser, whereas the gene conservation analysis was performed using UCSC Genome Browser (https://genome.ucsc.edu/). For enhancer analysis, the WashU Epigenome Browser (http://epigenomegateway.wustl.edu/browser/) (Li et al. 2019) was used, with tracks for RAMPAGE Seq data on NSC and neural cells.

### IDH-mutation Analysis

Raw RNAseq fastq files from initial and recurrent glioma pairs were downloaded from the European Genome-Phenome Archive under accession numbers EGAS00001001033 and EGAS00001001880, as well as the Genomic Data Commons Legacy Archive GBM and LGG datasets (https://portal.gdc.cancer.gov/legacy-archive) [PMID: 26373279, 28263318, 28697342]. Together, the samples used for these analyses comprise the publicly available expression data for the glioma pairs that are part of the Glioma Longitudinal Analysis (GLASS) Consortium (https://www.synapse.org/#!Synapse:syn17038081/). Each fastq file was preprocessed using fastp v0.20.0 and then input into kallisto v0.46.0 using Ensembl v75 non-coding RNA as the reference index. Transcript per million (TPM) values output by kallisto was used for downstream analyses. The sample sizes for each group are as follows: initial IDH mut (n = 10), initial IDH wt (n = 31), recurrent IDH mut (n = 10), and recurrent IDH wt (n = 31). Both the initial and recurrent tumors come from the same patients, so the numbers are the same for each IDHmut/IDHwt category.

### Hi-C

Hi-C contact matrices, loop calls, H3K27ac ChIP peaks, and CTCF ChIP peaks from primary glioblastoma cultures were accessed from GEO accession GSE121601 and https://wangftp.wustl.edu/hubs/johnston_gallo/ (Johnston et al. 2019). Two-dimensional DNA contact matrices were displayed using Juicebox 1.11.08 (Durand et al. 2016) at 5-kb resolution with balanced normalization and with identical color range applied to all panels. One-dimensional tracks were displayed using IGV (Robinson et al. 2011).

### RNA-Seq

RNA was extracted using RNeasy kit (Qiagen). For ribosomal depletion, 1–2 mg of total RNA was used for Ribo-Zero rRNA Removal kit (Illumina) according to the manufacturer’s protocol. Ribosomal-depleted library was processed for TruSeq Stranded Total RNA sequencing according to the manufacturer’s protocol (Illumina). The quality and quantity of total RNA and final libraries were assessed using Agilent Technologies 2100 Bioanalyzer. RIN (RNA Integrity Number) of >9 was used for the sequencing experiments.

### Data Analysis

For RNA-Sequencing data, reads were assessed for sequencing quality using FASTQC (v.0.11.7). Paired-end reads were aligned and counted using STAR (v.2.6.0c). STAR genome index was generated using GRCh38.p13 assembly and GRCh38.101 GTF from Ensembl. Raw counts were pre-filtered on the basis on row-wise sums across all samples with a cutoff of 5. Differential expression analysis was computed using DESeq2 (v1.28.1). The network map was generated using the BioBox Platform (https://biobox.io), a cloud genomic data analytics platform.

## DATA ACCESS

All raw and processed sequencing data generated in this study have been submitted to the NCBI Gene Expression Omnibus (GEO; https://www.ncbi.nlm.nih.gov/geo/) under accession number GSE157506.

## ACKNOWLEDGEMENTS

We would like to acknowledge the laboratory of Dr. Peter Dirks for providing the HFNS 6562, HFNS 7450, GliNS1, Gli432, and Gli489 cell lines, the lab of Dr. Michael Taylor for iPSCs, the lab of Dr. Phil Marsden for providing HUVECs, Dr. Mark Minden for the Jukrat, K562, and AML5 lines, and the lab of Dr. Brent Derry for DLD1 and HCT116 lines. We would also like to thank Dr. Paulo Amaral for his invaluable feedback. We would also like to acknowledge Emily Reddy from the Flow Cytometry Facility at Peter Gilgan Centre for Research and Learning for her help with the cell cycle analysis experiments.

## Notes

### Competing Interest Statement

The authors have declared no competing interest.

## REFERENCES

Agnihotri S, Burrell KE, Wolf A, Jalali S, Hawkins C, Rutka JT, Zadeh G. 2013. Glioblastoma, a brief review of history, molecular genetics, animal models and novel therapeutic strategies. Arch Immunol Ther Exp (Warsz) 61: 25–41.

Aldape K, Zadeh G, Mansouri S, Reifenberger G, von Deimling A. 2015. Glioblastoma: pathology, molecular mechanisms and markers. Acta Neuropathol 129: 829–848.

Bao S, Wu Q, McLendon RE, Hao Y, Shi Q, Hjelmeland AB, Dewhirst MW, Bigner DD, Rich JN. 2006. Glioma stem cells promote radioresistance by preferential activation of the DNA damage response. Nature 444: 756–760.

Beck B, Blanpain C. 2013. Unravelling cancer stem cell potential. Nat Rev Cancer 13: 727–738.

Bhan A, Soleimani M, Mandal SS. 2017. Long noncoding RNA and cancer: A new paradigm. Cancer Res 77: 3965–3981.

Bradner JE, Hnisz D, Young RA. 2017. Transcriptional Addiction in Cancer. Cell 168: 629–643.

Chang HC, Huang HC, Juan HF, Hsu CL. 2019. Investigating the role of super-enhancer RNAs underlying embryonic stem cell differentiation. BMC Genomics 20: 1–12.

Cheng JH, Pan DZC, Tsai ZTY, Tsai HK. 2015. Genome-wide analysis of enhancer RNA in gene regulation across 12 mouse tissues. Sci Rep 5: 1–9.

Chipumuro E, Marco E, Christensen CL, Kwiatkowski N, Zhang T, Hatheway CM, Abraham BJ, Sharma B, Yeung C, Altabef A, et al. 2014. CDK7 inhibition suppresses super-enhancer-linked oncogenic transcription in MYCN-driven cancer. Cell 159: 1126–1139.

Christensen CL, Kwiatkowski N, Abraham BJ, Carretero J, Al-Shahrour F, Zhang T, Chipumuro E, Herter-Sprie GS, Akbay EA, Altabef A, et al. 2014. Targeting Transcriptional Addictions in Small Cell Lung Cancer with a Covalent CDK7 Inhibitor. Cancer Cell 26: 909–922.

Das S, Srikanth M, Kessler J a. 2008. Cancer stem cells and glioma. Nat Clin Pract Neurol 4: 427–35.

Durand NC, Robinson JT, Shamim MS, Machol I, Mesirov JP, Lander ES, Aiden EL. 2016. Juicebox Provides a Visualization System for Hi-C Contact Maps with Unlimited Zoom. Cell Syst 3: 99–101.

Flicek P, Amode MR, Barrell D, Beal K, Billis K, Brent S, Carvalho-Silva D, Clapham P, Coates G, Fitzgerald S, et al. 2014. Ensembl 2014. Nucleic Acids Res 42: 749–755.

Flynn RA, Chang HY. 2014. Long noncoding RNAs in cell-fate programming and reprogramming. Cell Stem Cell 14: 752–761.

Gao X, He H, Zhu X, Xie S, Cao Y. 2019. LncRNA SNHG20 promotes tumorigenesis and cancer stemness in glioblastoma via activating PI3K/Akt/mTOR signaling pathway. Neoplasma 66: 532–542.

Gimple RC, Bhargava S, Dixit D, Rich JN. 2019. Glioblastoma stem cells: Lessons from the tumor hierarchy in a lethal cancer. Genes Dev 33: 591–609.

Guttman M, Donaghey J, Carey BW, Garber M, Grenier JK, Munson G, Young G, Lucas AB, Ach R, Bruhn L, et al. 2011. lincRNAs act in the circuitry controlling pluripotency and differentiation. Nature 477: 1–11.

Guttman M, Rinn JL. 2012. Modular regulatory principles of large non-coding RNAs. Nature 482: 339–46.

Hnisz D, Abraham BJ, Lee TI, Lau A, Saint-André V, Sigova AA, Hoke HA, Young RA. 2013. XSuper-enhancers in the control of cell identity and disease. Cell 155: 934.

Huang J, Li K, Cai W, Liu X, Zhang Y, Orkin SH, Xu J, Yuan GC. 2018. Dissecting super-enhancer hierarchy based on chromatin interactions. Nat Commun 9.

Jiang YY, Lin DC, Mayakonda A, Hazawa M, Ding LW, Chien WW, Xu L, Chen Y, Xiao JF, Senapedis W, et al. 2017. Targeting super-enhancer-Associated oncogenes in oesophageal squamous cell carcinoma. Gut 66: 1358–1368.

Johnston MJ, Nikolic A, Ninkovic N, Guilhamon P, Cavalli FMG, Seaman S, Zemp FJ, Lee J, Abdelkareem A, Ellestad K, et al. 2019. High-resolution structural genomics reveals new therapeutic vulnerabilities in glioblastoma. Genome Res 29: 1211–1222.

Khalil AM, Guttman M, Huarte M, Garber M, Raj A, Rivea Morales D, Thomas K, Presser A, Bernstein BE, van Oudenaarden A, et al. 2009. Many human large intergenic noncoding RNAs associate with chromatin-modifying complexes and affect gene expression. Proc Natl Acad Sci U S A 106: 11667–11672.

Kopp F, Mendell JT. 2018. Functional Classification and Experimental Dissection of Long Noncoding RNAs. Cell 172: 393–407.

Kuşoğlu A, Biray Avcı Ç. 2019. Cancer stem cells: A brief review of the current status. Gene 681: 80–85.

Lai F, Damle SS, Ling KK, Rigo F. 2020. Directed RNase H Cleavage of Nascent Transcripts Causes Transcription Termination. Mol Cell 77: 1032–1043.e4.

Lam MTY, Li W, Rosenfeld MG, Glass CK. 2014. Enhancer RNAs and regulated transcriptional programs. Trends Biochem Sci 39: 170–182.

Lan X, Jörg DJ, Cavalli FMG, Richards LM, Nguyen L V., Vanner RJ, Guilhamon P, Lee L, Kushida MM, Pellacani D, et al. 2017. Fate mapping of human glioblastoma reveals an invariant stem cell hierarchy. Nature 549: 227–232.

Lathia J, Mack S. 2015. Cancer stem cells in glioblastoma. Genes … 1203–1217.

Lee JS, Mendell JT. 2020. Antisense-Mediated Transcript Knockdown Triggers Premature Transcription Termination. Mol Cell 77: 1044–1054.e3.

Lennox KA, Behlke MA. 2016. Cellular localization of long non-coding RNAs affects silencing by RNAi more than by antisense oligonucleotides. Nucleic Acids Res 44: 863–877.

Li D, Hsu S, Purushotham D, Sears RL, Wang T. 2019. WashU Epigenome Browser update 2019. Nucleic Acids Res 47: W158–W165.

Li W, Notani D, Rosenfeld MG. 2016. Enhancers as non-coding RNA transcription units: Recent insights and future perspectives. Nat Rev Genet 17: 207–223.

Li Y, Rivera CM, Ishii H, Jin F, Selvaraj S, Lee AY, Dixon JR, Ren B. 2014. CRISPR reveals a distal super-enhancer required for Sox2 expression in mouse embryonic stem cells. PLoS One 9: 1–17.

Lin CY, Erkek S, Tong Y, Yin L, Federation AJ, Zapatka M, Haldipur P, Kawauchi D, Risch T, Warnatz HJ, et al. 2016. Active medulloblastoma enhancers reveal subgroup-specific cellular origins. Nature 530: 57–62.

Lu C, Ward PS, Kapoor GS, Rohle D, Turcan S, Abdel-Wahab O, Edwards CR, Khanin R, Figueroa ME, Melnick A, et al. 2012. IDH mutation impairs histone demethylation and results in a block to cell differentiation. Nature 483: 474–478.

Ma Q, Long W, Xing C, Chu J, Luo M, Wang HY, Liu Q, Wang RF. 2018. Cancer Stem Cells and Immunosuppressive Microenvironment in Glioma. Front Immunol 9: 2924.

Mineo M, Ricklefs F, Rooj AK, Lyons SM, Ivanov P, Ansari KI, Nakano I, Chiocca EA, Godlewski J, Bronisz A. 2016. The long non-coding RNA-HIF1A-AS2 facilitates the maintenance of mesenchymal glioblastoma stem-like cells in hypoxic niches. Cell Rep 15: 2500–2509.

Nassar D, Blanpain C. 2016. Cancer Stem Cells: Basic Concepts and Therapeutic Implications. Annu Rev Pathol Mech Dis 11: 47–76.

Nørøxe DS, Poulsen HS, Lassen U. 2016. Hallmarks of glioblastoma:a systematic review. ESMO Open 1.

Ooi WF, Xing M, Xu C, Yao X, Ramlee MK, Lim MC, Cao F, Lim K, Babu D, Poon LF, et al. 2016. Epigenomic profiling of primary gastric adenocarcinoma reveals super-enhancer heterogeneity. Nat Commun 7: 1–17.

Ostrom QT, Cioffi G, Gittleman H, Patil N, Waite K, Kruchko C, Barnholtz-Sloan JS. 2019. CBTRUS Statistical Report: Primary Brain and Other Central Nervous System Tumors Diagnosed in the United States in 2012-2016. Neuro Oncol 21: V1–V100.

Ounzain S, Micheletti R, Arnan C, Plaisance I, Cecchi D, Schroen B, Reverter F, Alexanian M, Gonzales C, Ng SY, et al. 2015. CARMEN, a human super enhancer-associated long noncoding RNA controlling cardiac specification, differentiation and homeostasis. J Mol Cell Cardiol 89: 98–112.

Plaks V, Kong N, Werb Z. 2015. The cancer stem cell niche: How essential is the niche in regulating stemness of tumor cells? Cell Stem Cell 16: 225–238.

Pott S, Lieb JD. 2015. What are super-enhancers? Nat Genet 47: 8–12.

Rinaldi C, Wood MJA. 2018. Antisense oligonucleotides: The next frontier for treatment of neurological disorders. Nat Rev Neurol 14: 9–22.

Rinn JL, Chang HY. 2012. Genome regulation by long noncoding RNAs. Ann Rev Biochem 81: 145–166.

Robinson JT, Thorvaldsdóttir H, Winckler W, Guttman M, Lander ES, Getz G, Mesirov JP. 2011. Integrative Genome Viewer. Nat Biotechnol 29: 24–6.

Sampetrean O, Saya H. 2013. Characteristics of glioma stem cells. Brain Tumor Pathol 30: 209–214.

Sanai N, varez-Buylla A, Berger MS. 2005. Neural stem cells and the origin of gliomas. N Engl J Med 353: 811–822.

Schmitt AM, Chang HY. 2016. Long Noncoding RNAs in Cancer Pathways. Cancer Cell 29.

Sengupta S, George RE. 2017. Super-Enhancer-Driven Transcriptional Dependencies in Cancer. Trends in Cancer 3: 269–281.

Singh SK, Hawkins C, Clarke ID, Squire JA, Bayani J, Hide T, Henkelman RM, Cusimano MD, Dirks PB. 2004. Identification of human brain tumour initiating cells. Nature 432: 396–401.

Stupp R, Mason WP, van den Bent MJ, Weller M, Fisher B, Taphoorn MJB, Belanger K, Brandes AA, Marosi C, Bogdahn U, et al. 2005. Radiotherapy plus Concomitant and Adjuvant Temozolomide for Glioblastoma. N Engl J Med 352: 987–996.

Sur I, Taipale J. 2016. The role of enhancers in cancer. Nat Rev Cancer 16: 483–493.

Suvà ML, Rheinbay E, Gillespie SM, Patel AP, Wakimoto H, Rabkin SD, Riggi N, Chi AS, Cahill DP, Nahed B V., et al. 2014. Reconstructing and reprogramming the tumor-propagating potential of glioblastoma stem-like cells. Cell 157: 580–594.

Suvà ML, Tirosh I. 2020. The Glioma Stem Cell Model in the Era of Single-Cell Genomics. Cancer Cell 37: 630–636.

Takahashi K, Yamanaka S. 2006. Induction of Pluripotent Stem Cells from Mouse Embryonic and Adult Fibroblast Cultures by Defined Factors. Cell 126: 663–676.

Thandapani P. 2019. Super-enhancers in cancer. Pharmacol Ther 199: 129–138.

Turcan S, Makarov V, Taranda J, Wang Y, Fabius AWM, Wu W, Zheng Y, El-Amine N, Haddock S, Nanjangud G, et al. 2018. Mutant-IDH1-dependent chromatin state reprogramming, reversibility, and persistence. Nat Genet 50: 62–72.

Ulitsky I, Bartel DP. 2013. LincRNAs: Genomics, evolution, and mechanisms. Cell 154: 26–46.

Venere M, Fine HA, Dirks PB, Rich JN. 2011. Cancer stem cells in gliomas: Identifying and understanding the apex cell in cancer’s hierarchy. Glia 59: 1148–1154.

Wang Y, Xu Z, Jiang J, Xu C, Kang J, Xiao L, Wu M, Xiong J, Guo X, Liu H. 2013. Endogenous miRNA Sponge lincRNA-RoR Regulates Oct4, Nanog, and Sox2 in Human Embryonic Stem Cell Self-Renewal. Dev Cell 25: 69–80.

Wang Y, Zhang T, Kwiatkowski N, Abraham BJ, Lee TI, Xie S, Yuzugullu H, Von T, Li H, Lin Z, et al. 2015. CDK7-Dependent Transcriptional Addiction in Triple-Negative Breast Cancer. Cell 163: 174–186.

Weintraub AS, Li CH, Zamudio A V., Sigova AA, Hannett NM, Day DS, Abraham BJ, Cohen MA, Nabet B, Buckley DL, et al. 2017. YY1 Is a Structural Regulator of Enhancer-Promoter Loops. Cell 171: 1573–1588.e28.

Whyte WA, Orlando DA, Hnisz D, Abraham BJ, Lin CY, Kagey MH, Rahl PB, Lee TI, Young RA. 2013. Master transcription factors and mediator establish super-enhancers at key cell identity genes. Cell 153: 307–319.

Xiang JF, Yin QF, Chen T, Zhang Y, Zhang XO, Wu Z, Zhang S, Wang H Bin, Ge J, Lu X, et al. 2014. Human colorectal cancer-specific CCAT1-L lncRNA regulates long-range chromatin interactions at the MYC locus. Cell Res 24: 513–531.

Xie JJ, Jiang YY, Jiang Y, Li CQ, Lim MC, An O, Mayakonda A, Ding LW, Long L, Sun C, et al. 2018. Super-Enhancer-Driven Long Non-Coding RNA LINC01503, Regulated by TP63, Is Over-Expressed and Oncogenic in Squamous Cell Carcinoma. Gastroenterology 154: 2137–2151.e1.

Zhou B, Wang L, Zhang S, Bennett BD, He F, Zhang Y, Xiong C, Han L, Diao L, Li P, et al. 2016. INO80 governs superenhancer-mediated oncogenic transcription and tumor growth in melanoma. Genes Dev 30: 1440–1453.

