## Supplemental Figure 1 for "*CASCADES*, a novel *SOX2* super-enhancer associated long noncoding RNA, regulates cancer stem cell specification and differentiation in glioblastoma multiforme"

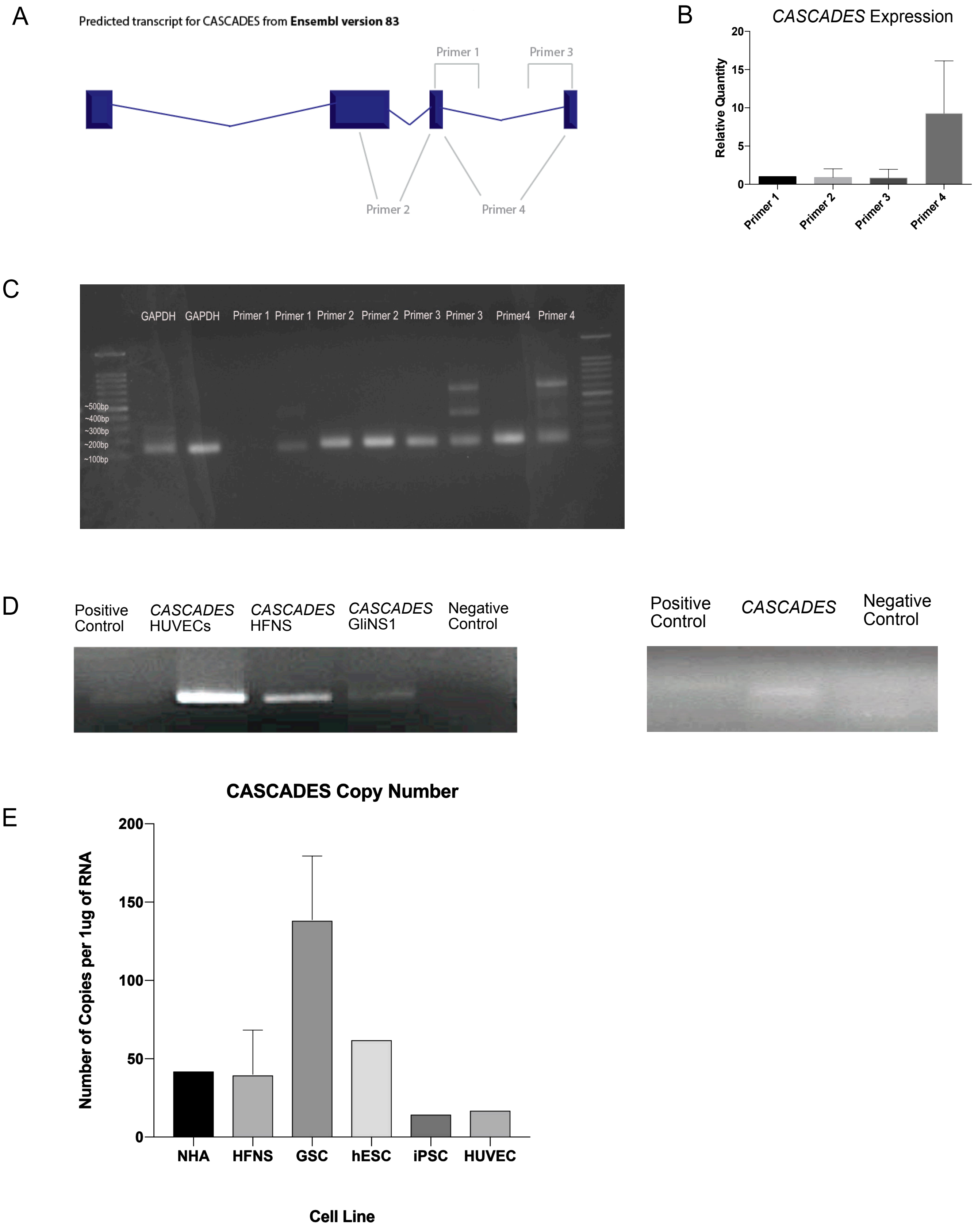


**Supplementary Figure 1. A, B.** The expression of *CASCADES* upon primer crawling designed across different areas of the first transcript. **C.** 3'Rapid Amplification of cDNA Ends (RACE) of *CASCADES* in glioma stem cells (GliNS1), normal human fetal neural stem cells (HFNS), and human umbilical vein endothelial cells (HUVECs). **D.** 5'RACE of *CASCADES* in glioma stem cells (GliNS1) cell line. **E.** The copy number of *CASCADES* across different cell lines.
