## Supplemental Figure 2 for "*CASCADES*, a novel *SOX2* super-enhancer associated long noncoding RNA, regulates cancer stem cell specification and differentiation in glioblastoma multiforme"

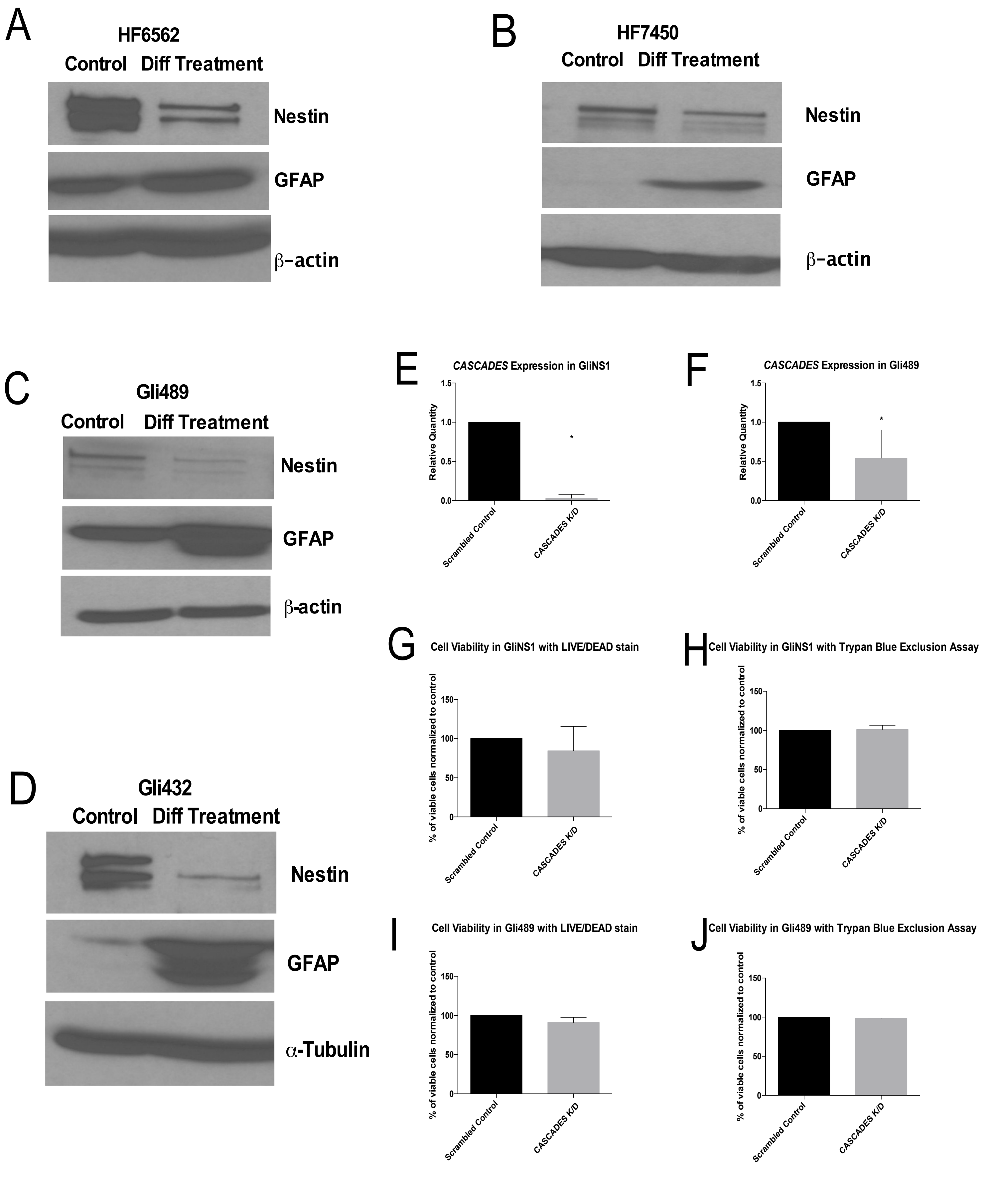


**Supplementary Figure 2.**

Two HFNS cell lines, HF6562 (A) and HF7450 (B) as well as two GSC lines, Gli489 (C) and Gli432 (D) were forced to differentiate into astrocytes. The expression of neural stem cell marker, Nestin had decreased upon forced differentiation, while expression of glial fibrillary acidic protein (GFAP) increased.

The *CASCADES* expression was knockdown using siRNA designed against the *CASCADES* transcript. The expression of *CASCADES* was significantly decreased in both GliNS1 (E) and Gli489 (F) cell lines. Furthermore, the knockdown of *CASCADES* did not adversely affect cell viability in either GliNS1 (G,H), or Gli489 (I, J), as assessed by both LIVE/DEAD staining and Trypan Blue Exclusion assay.
