## Supplemental Figure 3 for "*CASCADES*, a novel *SOX2* super-enhancer associated long noncoding RNA, regulates cancer stem cell specification and differentiation in glioblastoma multiforme"

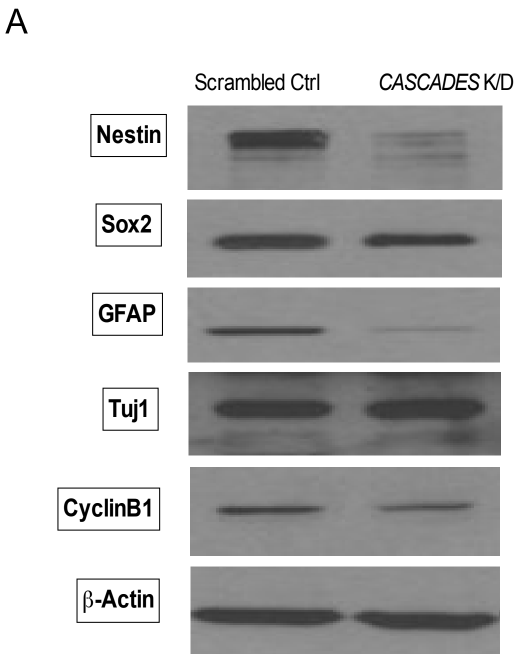


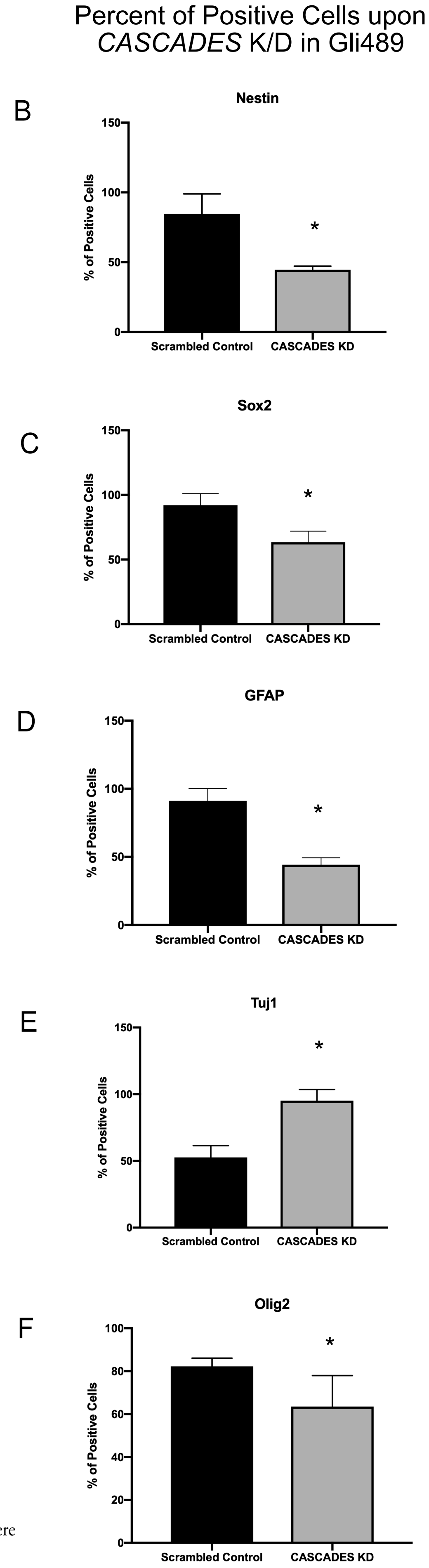


**Supplementary Figure 3.** The glioma stem cell line Gli489 (N=3/group) was treated with siRNA directed against *CASCADES* or universal negative control (scrambled) for 72 hours.
**A.** The Western blotting analysis of various neural stemness markers in *CASCADES* K/D vs. scrambled control
**B-F.** The percentage of positive cells out of total cells identified through confocal microscopy were plotted as bar graphs.
