## Supplemental Figure 4 for "*CASCADES*, a novel *SOX2* super-enhancer associated long noncoding RNA, regulates cancer stem cell specification and differentiation in glioblastoma multiforme"

### *CASCADES* Knockdown in GliNS1

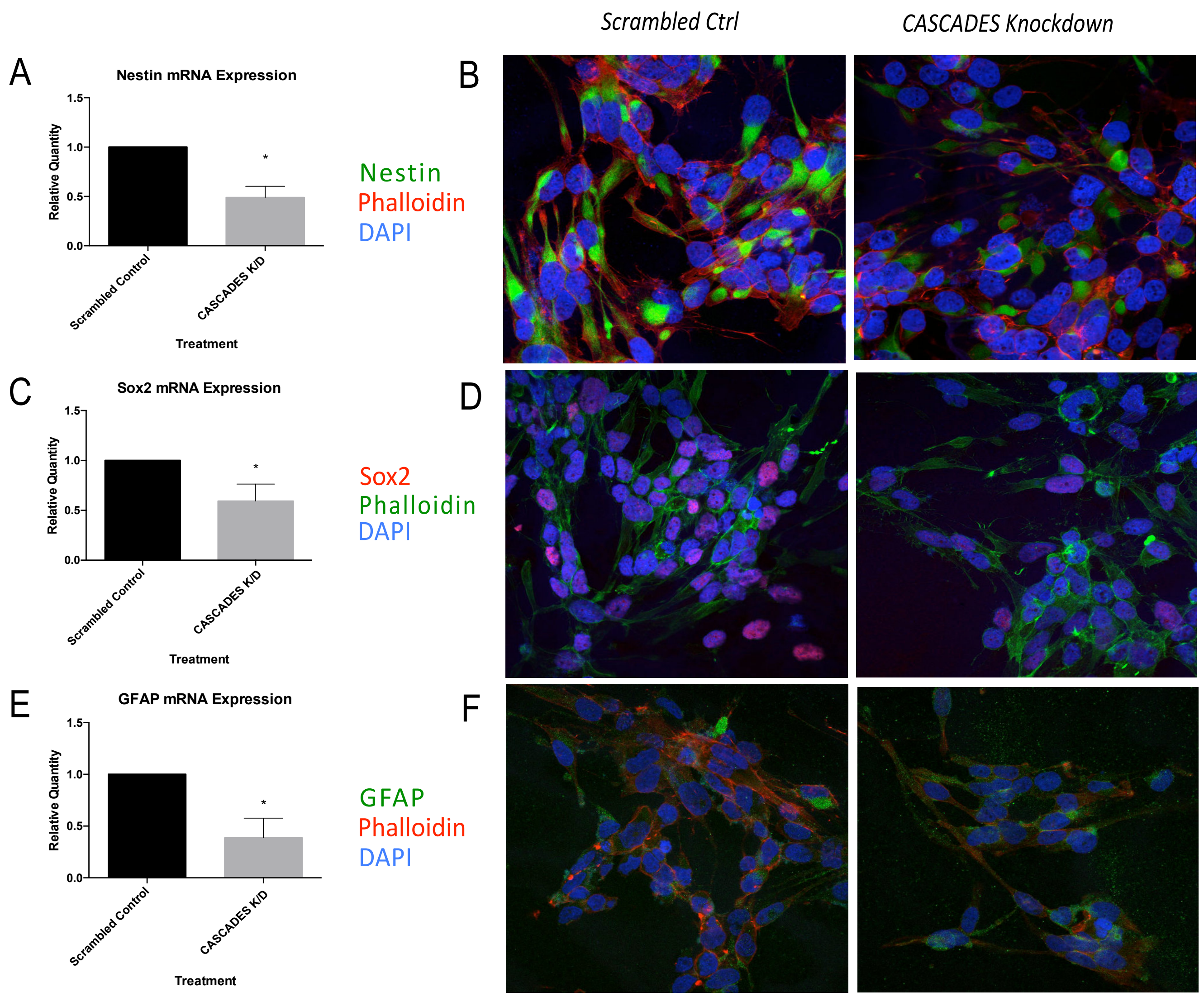


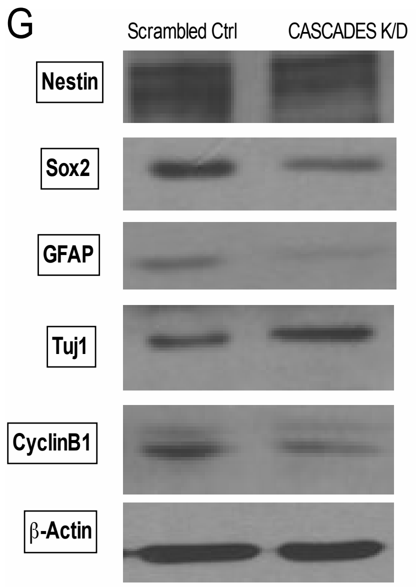


**Supplementary Figure 4.** The glioma stem cell line GliNS1 (N=3/group) was treated with siRNA directed against *CASCADES* or universal negative control (scrambled) for 72 hours.
The stemnss markers, Nestin (A,B,&G) and Sox2 (C,D,&G) were significant decreased after *CASCADES* knockdown, compared to scrambled controls. Furthermore, the expression of GFAP, an astrocyte marker, also showed a decrease (E,F,&G). In comparison, the differentiated neuron marker, Tuj1 was greatly increased in expression, after *CASCADES* knockdown. Furthermore, the knockdown of *CASCADES* decreased the expression of cyclin B1 (G), compared to scrambled controls.
