## Supplemental Figure 6 for "*CASCADES*, a novel *SOX2* super-enhancer associated long noncoding RNA, regulates cancer stem cell specification and differentiation in glioblastoma multiforme"

### *CASCADES* Knockdown in HFNS 6562

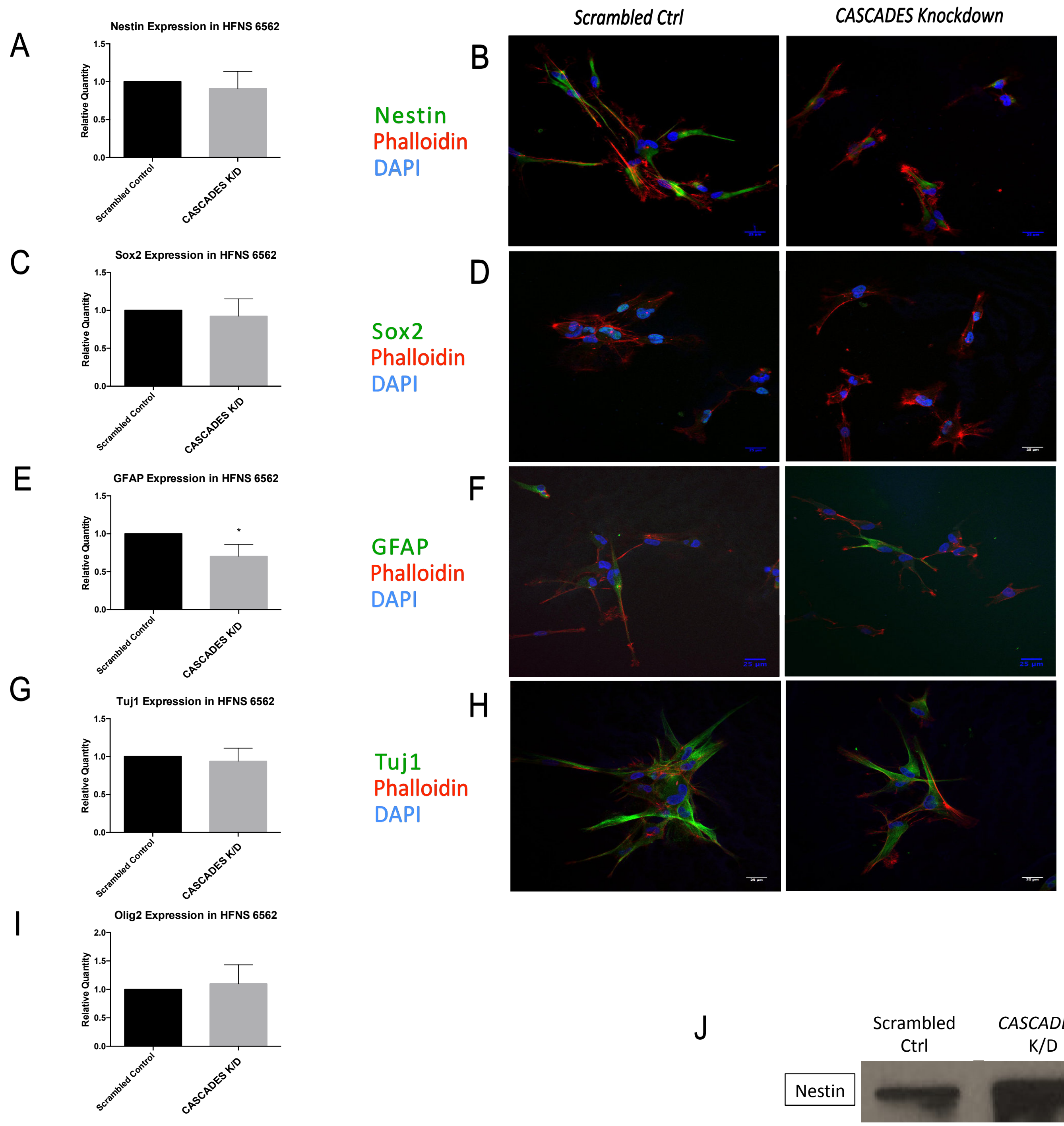


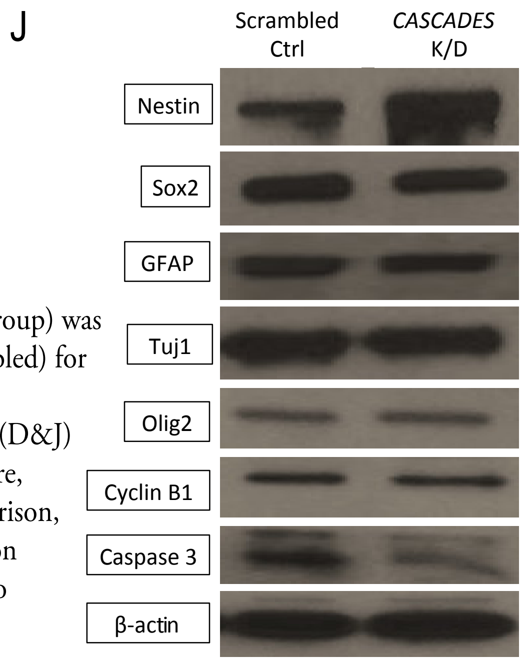


**Supplementary Figure 6.** The human fetal neural stem cell line HFNS 6562 (N=3/group) was treated with siRNA targeting against *CASCADES* or universal negative control (scrambled) for 72 hours.

The stemnss marker, Nestin (B&J) showed slight increase in expression, whereas Sox2 (D&J) decreased after *CASCADES* knockdown, compared to scrambled controls. Furthermore, the expression of GFAP, an astrocyte marker, also showed a decrease (F&J). In comparison, Tuj1 (H&J), Olig2 (J), and cyclin B1 (J), did not show significantly different expression after *CASCADES* knockdown when compared to scrambled controls. Furthermore, no statistically significant differences were detected at the mRNA level (A,C,E,&G).
