## Supplemental Figure 7 for "*CASCADES*, a novel *SOX2* super-enhancer associated long noncoding RNA, regulates cancer stem cell specification and differentiation in glioblastoma multiforme"

### *CASCADES* Localization Controls

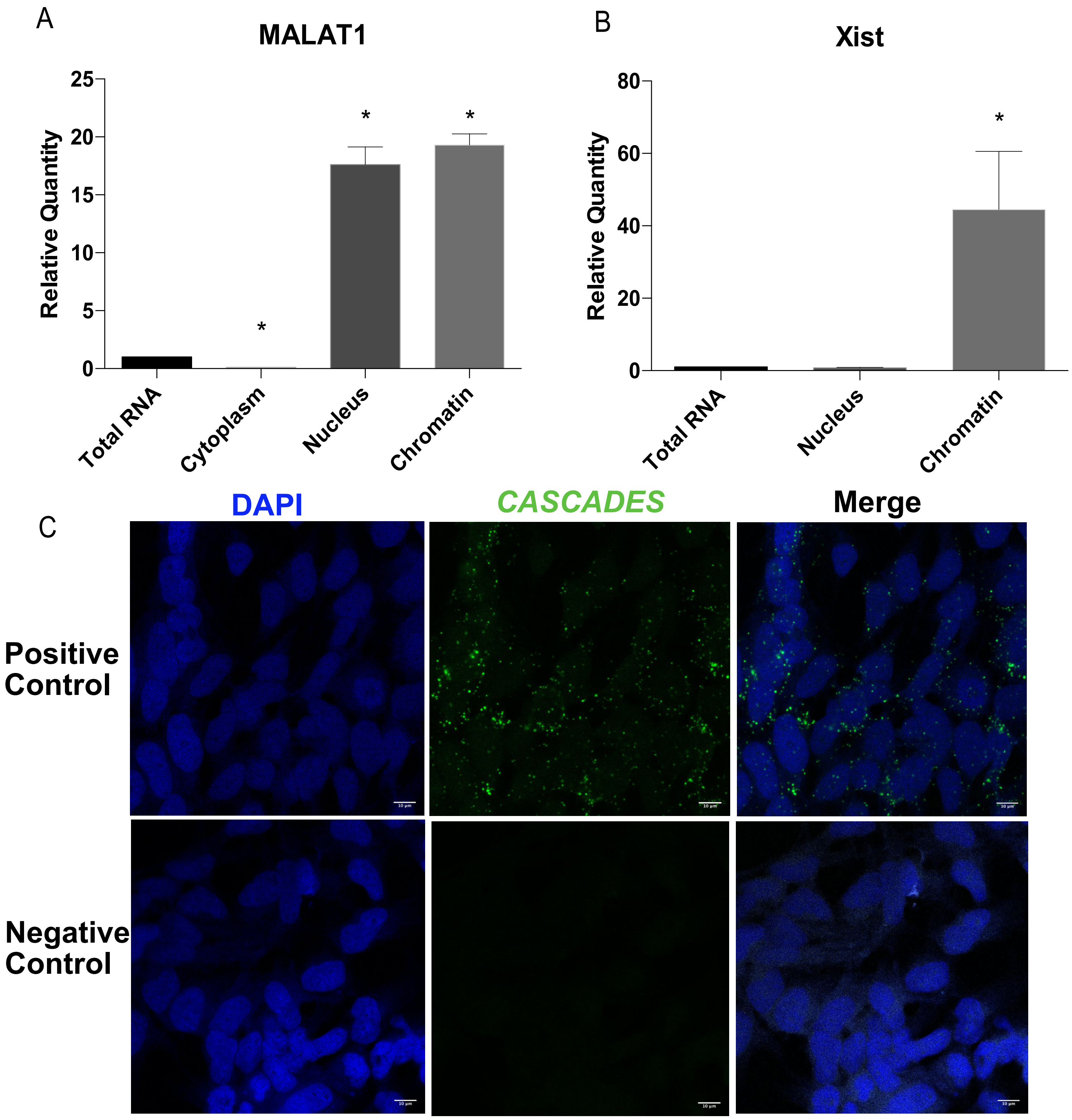


**Supplementary Figure 7.** The *CASCADES* expression was assessed in different cellular fractions and compared to total RNA.
(A) MALAT1 (B) and Xist served as controls.
(C) The positive and negative controls for the RNA *in situ* hybridization.
