## Supplemental Figure 8 for "*CASCADES*, a novel *SOX2* super-enhancer associated long noncoding RNA, regulates cancer stem cell specification and differentiation in glioblastoma multiforme"

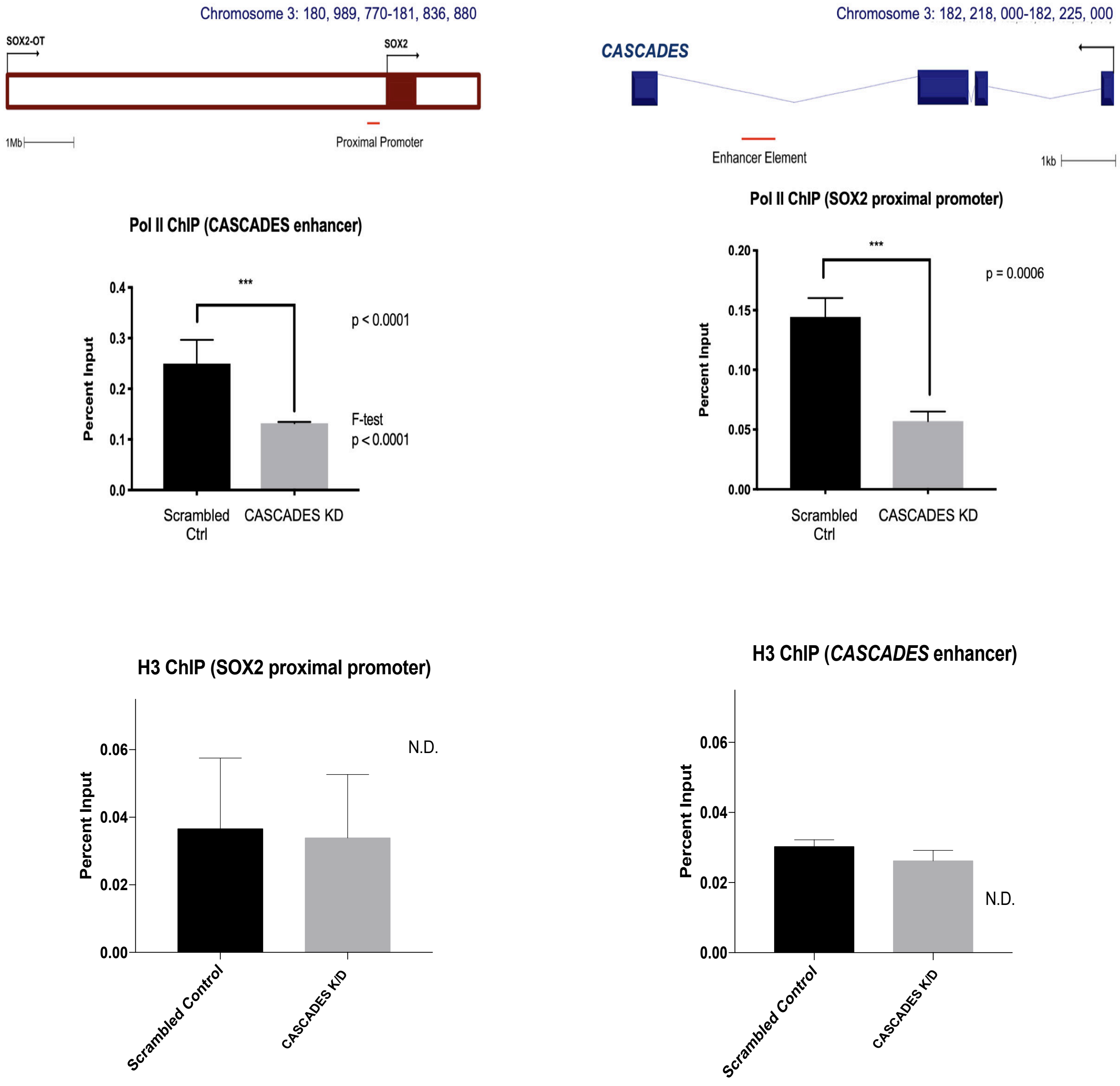


**Supplementary Figure 8.** Chromatin immunoprecipitation was performed for Rad21, YY1, and RNA Pol II. H3 served as control.
