## Supplemental Figure 9 for "*CASCADES*, a novel *SOX2* super-enhancer associated long noncoding RNA, regulates cancer stem cell specification and differentiation in glioblastoma multiforme"

### *CASCADES* Knockdown using ASO

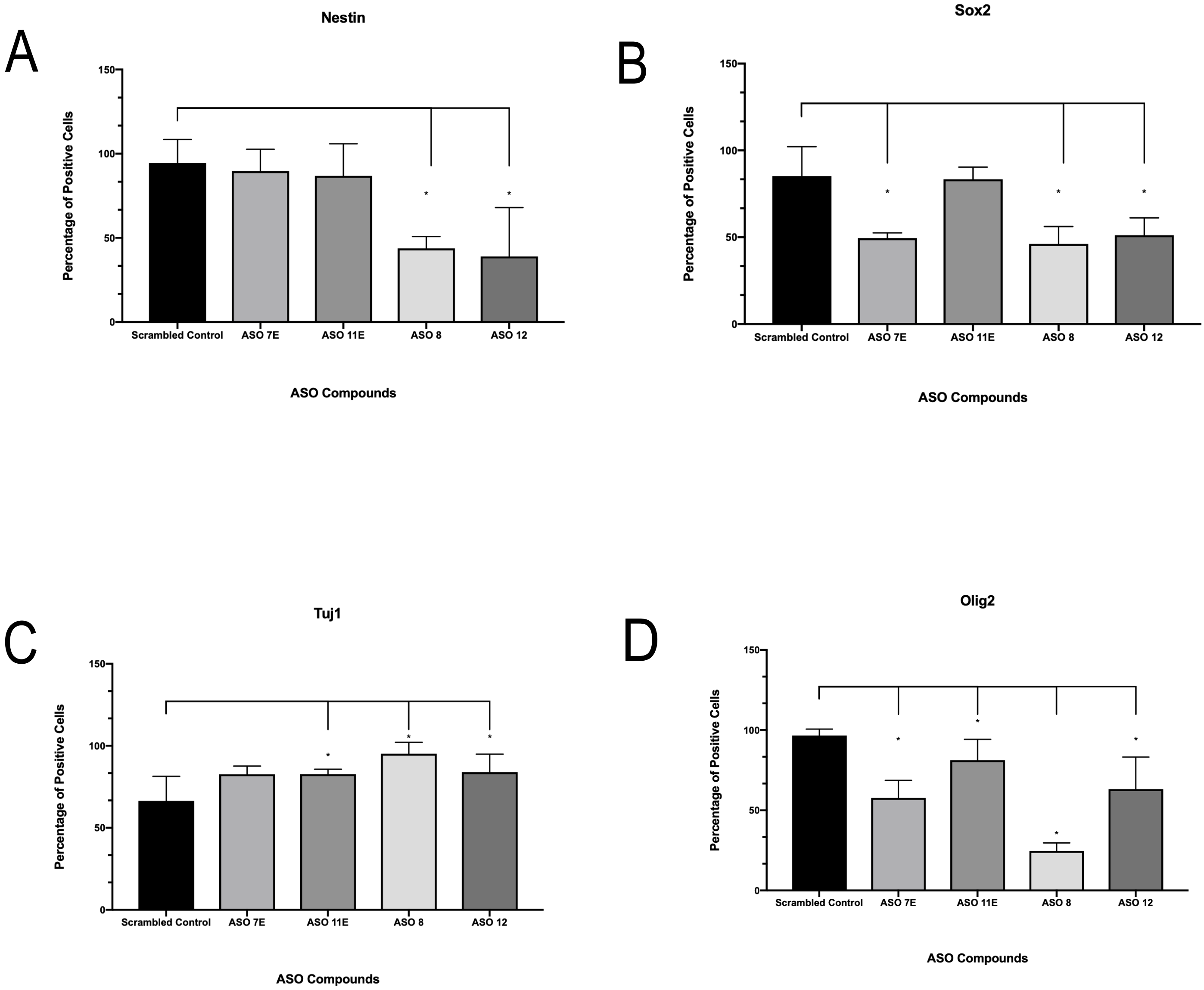


**Supplementary Figure 9.** The *CASCADES* expression was knocked down using antisense oligonucleotides designed against the *CASCADES* transcript or the enhancer element found within the transcript.
The knockdown of *CASCADES* using the ASO 8 & ASO 12 resulted in significant reduction of percentage of cells expressing Nestin (A), Sox2(B), and Olig2 (D), and an increase in Tuj1 (C).
