## Supplemental Figure 10 for "*CASCADES*, a novel *SOX2* super-enhancer associated long noncoding RNA, regulates cancer stem cell specification and differentiation in glioblastoma multiforme"

### *CASCADES* Knockdown

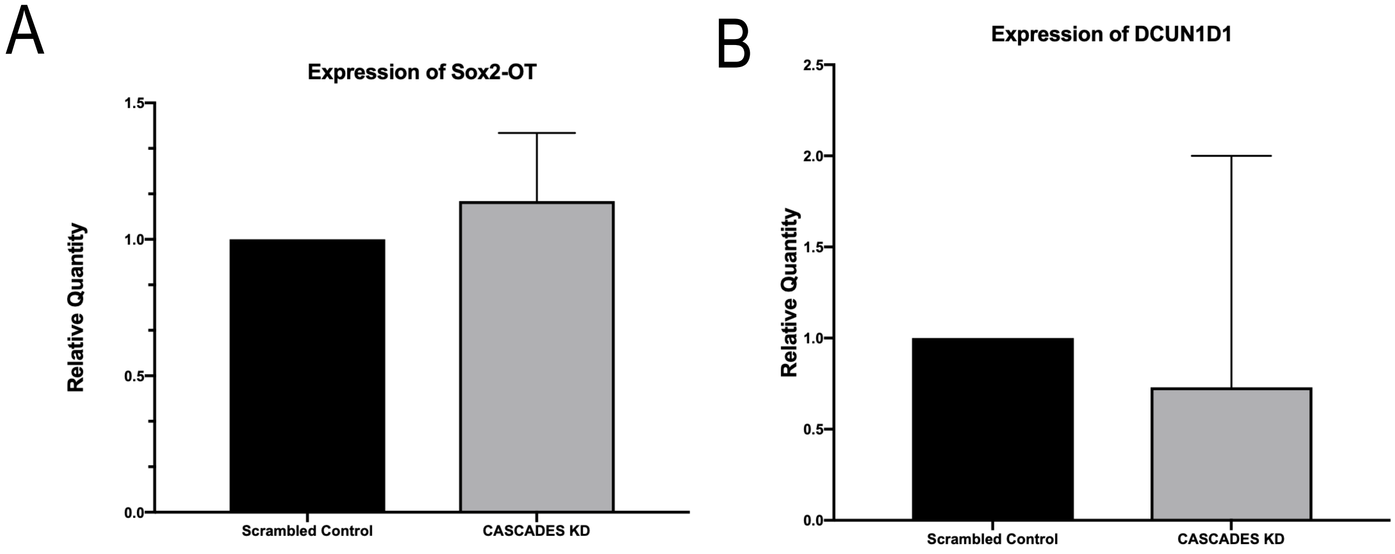


**Supplementary Figure 10.** The knockdown of *CASCADES* using the siRNA did not significantly affect the neighboring genes, Sox2-OT (A) and DCUN1D1 (B).
