## Supplemental Table 1 for "*CASCADES*, a novel *SOX2* super-enhancer associated long noncoding RNA, regulates cancer stem cell specification and differentiation in glioblastoma multiforme"

| Gene Symbol | Nearby Gene | Strand | Location | Splice Variants | Longest Transcript Length |
|---|---|---|---|---|---|
| **XXbac-BPG27H4.8** | Oct3/Oct4(POU5F1) | Reverse | chrom 6 | 1 | 799 |
| **HCG20** | Oct3/Oct4(POU5F1) | Forward | chrom 6 | 2 | 1202 |
| **XXbac-BPG181B23.7** | Oct3/Oct4(POU5F1) | Reverse | chrom 6 | 1 | 1207 |
| **LINC01149** | Oct3/Oct4(POU5F1) | Forward | chrom 6 | 1 | 2141 |
| **RP11-139K4.2** | Sox2 | Reverse | chrom 3 | 1 | 621 |
| **RP11-416O18.2** | Sox2 | Forward | chrom 3 | 1 | 477 |
| **CASC11** | Myc & POU5F1B | Reverse | chrom 8 | 3 | 3301 |
| **CASC19** | Myc & POU5F1B | Reverse | chrom 8 | 1 | 367 |
| **PCAT1** | Myc & POU5F1B | Forward | chrom 8 | 2 | 1992 |
| **RP11-761N21.1** | Beta-Catenin (CTNNB1) | Forward | chrom 3 | 1 | 500 |
| **RP11-391M1.4** | Beta-Catenin (CTNNB1) | Forward | chrom 3 | 1 | 2122 |
| **LINC01060** | Rex1 (Zfp42) | Forward | chrom 4 | 6 | 2435 |
| **AE000658.31** | Sall2 | Reverse | chrom 14 | 1 | 570 |
| **RP11-998D10.7** | Sall2 | Forward | chrom 14 | 1 | 419 |
| **AL161668.5** | Sall2 | Forward | chrom 14 | 2 | 478 |
| **RP11-219E7.4** | Sall2 | Reverse | chrom 14 | 1 | 351 |
| **RP11-219E7.3** | Sall2 | Reverse | chrom 14 | 2 | 549 |
| **LINC00945** | Olig1/Olig2 | Forward | chrom 21 | 1 | 753 |
| **AP000289.6** | Olig1/Olig2 | Reverse | chrom 21 | 1 | 1361 |
| **AP000569.9** | Olig1/Olig2 | Reverse | chrom 21 | 1 | 591 |

**Supplementary Table 1:** Top 20 lncRNA targets.
